# Long Non-Coding RNA Generated from *CDKN1A* Gene by Alternative Polyadenylation Regulates p21 Expression during DNA Damage Response

**DOI:** 10.1101/2023.01.10.523318

**Authors:** Michael R. Murphy, Anthony Ramadei, Ahmet Doymaz, Sophia Varriano, Devorah Natelson, Amy Yu, Sera Aktas, Marie Mazzeo, Michael Mazzeo, George Zakusilo, Frida E. Kleiman

**Author notes:** Corresponding author: FEK.

## Abstract

Alternative Polyadenylation (APA) is an emerging mechanism for dynamic changes in gene expression. Previously, we described widespread APA occurrence in introns during the DNA damage response (DDR). Here, we show that a DNA damage activated APA event occurs in the first intron of *CDKN1A*, inducing an alternate last exon (ALE)-containing lncRNA. We named this lncRNA SPUD (Selective Polyadenylation Upon Damage). SPUD localizes to polysomes in the cytoplasm and is detectable as multiple isoforms in available high throughput studies. SPUD has low abundance compared to the CDKN1A full-length isoform and is induced in cancer and normal cells under a variety of DNA damaging conditions in part through p53 transcriptional activation. RNA binding protein (RBP) HuR and the transcriptional repressor CTCF regulate SPUD levels. SPUD induction increases p21 protein, but not CDKN1A full-length levels, affecting p21 functions in cell-cycle, CDK2 expression, and cell viability. Like CDKN1A full-length isoform, SPUD can bind two competitive p21 translational regulators, the inhibitor calreticulin and the activator CUGBP1; SPUD can change their association with CDKN1A full-length in a DDR-dependent manner. Together, these results show a new regulatory mechanism by which a lncRNA controls p21 expression post-transcriptionally, highlighting lncRNA relevance in DDR progression and cellcycle.

## Introduction

The annotated human genome contains almost 200,000 transcripts from nearly 60,000 genes(1,2). Many of these additional transcripts are expressed from protein-coding genes as a product of alternative RNA processing, including splicing and polyadenylation (3,4). Alternate transcripts produced from the same protein-coding gene have the potential to be intimately linked in the regulation of the resultant protein by altering mRNA composition or generating distinct transcripts, while utilizing the same transcription and RNA processing machineries (5). One example are long non-coding RNAs (lncRNAs), which are characterized by poor conservation, low abundance (relative to mRNAs), and their induction in specific settings, such as certain tissues or in response to stress (6,7,8). The transient nature of lncRNA induction facilitates the fine-tuning of different cellular responses, such as the type and length of DNA damage response (DDR) (9,10).

Several lncRNAs have been described in close proximity to *CDKN1A* (11), the gene that codes for p21, that function either *in cis* affecting *CDKN1A* transcription or *in trans* by binding to other DDR factors (12,13,14,15). The cyclin-dependent kinase (CDK) inhibitor p21 has a variety of functions in cell-cycle regulation and DDR, playing a role in cell fate decision between senescence and apoptosis after DNA damage (reviewed in 16). Thus, *CDKN1A*, and perhaps other cell-cycle genes, exist as hubs for non-coding transcription to regulate cellular functions, such as DDR and cell-cycle progression, in addition to canonical protein production.

*CDKN1A* is a highly regulated transcriptional target of p53. A delayed in CDKN1A mRNA induction and p21 expression has been described for various stresses, such as UVC and during S-phase block with hydroxyurea (HU), despite comparable levels of RNA polymerase II and p53 at *CDKN1A* promoter (17,18). The delay in CDKN1A full-length mRNA induction is in part due to a block in transcription elongation somewhere in intron 1 (19,20,21,22). Whether the truncated CDKN1A transcript is processed or degraded has not currently been addressed.

Alternative polyadenylation (APA) is an effective regulator of cellular homeostasis (23,24,25). Approximately 20% of human genes have an intronic polyadenylation signal (PAS) with many tied to an alternate splicing event (alternate last exon; ALE, 26). A considerable number of intron-APA events occur in cancer cells that affect the coding region, generating in same cases either truncated and/or alternative C-termini protein products (27,28,29). Interestingly, intron-APA can also generate non-coding RNAs regulating its own protein-coding mRNA in response to UV damage (5). Our lab previously described an UV-induced increase in promoter-proximal APA events, specifically within introns (intron-APA), mediated by U1 snRNA (30–31) that are biased to genes with functions in DDR and cancer, including *CDKN1A* (32). It is not known whether these transcripts represent functional products of DDR or passive units generated during the suppression of transcription/processing of canonical mRNA; however, the high representation of intron-APA transcripts in RNAseq datasets and the presence of strong PAS sites in promoter-proximal introns suggests that these transcripts are indeed functional (33).

Here we provide insights into the truncated CDKN1A transcript generated by intron-APA after UV-treatment, which we named Selective Polyadenylation Upon DNA Damage (SPUD, 32). Our results indicate that SPUD is an ALE-containing lncRNA regulated by p53, which localizes to polysomes and is detectable in different datasets. SPUD has low abundance compared to CDKN1A full-length isoform and is induced in cancer and normal cells under a variety of damaging conditions. SPUD can functionally interact with different RBPs, including the p21 translational regulators calreticulin (CRT) and CUGBP1 consistent with SPUD role in regulating p21 expression and functions in cell-cycle, CDK2 expression, and cell viability. Together, our study reveals a new mechanism by which lncRNA SPUD regulates p21 expression at the translational level, highlighting SPUD relevance in cellular functions such as DDR progression and cell-cycle.

## Materials and Methods

### Cell lines, plasmids, and treatments

HCT116, HCT116 p21-/-, HCT116 p53 -/-, BJ, MCF7, and MDA-MB-231 cells were grown as previously described (34). HCT116 p21-/- cells were generously provided by Dr. Bert Vogelstein (Johns Hopkins University, Baltimore). APA isoforms were cloned into mammalian p3xFLAG-CMV10 and bacterial pET-42a(+) vectors. Mammalian vectors were treated with endotoxin removal component of Qiagen kit before transfections. UV treatment (20 Jm^2^) were as in (34). 1.7 mM hydroxyurea (HU, Sigma-Aldrich), 4 mM caffeine, or 10 μM etoposide (MilliporeSigma) were added directly to media before harvesting, with 16 h pretreatment for HU and etoposide and 1 h for caffeine (22). CUGBP1 and CRT expressing constructs were generously provided by Dr. Wilusz (University of Colorado) and Dr. Michalak (University of Alberta), respectively.

### Plasmid mutagenesis

CDKN1A intron 1 APA 3’ splice site was deleted using Q5 Site-Directed Mutagenesis Kit (New England Biolabs) as per the manufacturer’s instructions using forward primer (5’-TCCCCACCCCAAAATGACGCGCAGCC-3’) and reverse primer (5’-GGGGGAGAATGGGAGGGG-3’).

### RNA extraction and qRT-PCR assays

Total RNA purification and qRT-PCR were performed as described (32). RNA from nuclear and cytoplasmic compartments was extracted essentially as described (32). Relative RNA levels were calculated using 2-ΔΔCT method (36) and normalized against either Ubiquitin C (UBC) or β-Actin. Additional primers are described in Supplemental Figure S5.

### Knockdown expression

siRNA specific for human HuR, human CTCF, CRT, CUGBP1, a custom-made siRNA for SPUD and a control siRNA were obtained from Dharmacon RNA technologies. Knockdowns were done according to manufacturer’s protocol (Invitrogen) and tested by qRT-PCR and Western blot. SPUD knockdowns we tested by qRT-PCR.

### RNA stability assay

RNA was analyzed by qRT-PCR from cells treated with UV irradiation (20 Jm^2^), allowed to recover for 2 h and then treated with 2 μg/ml actinomycin D into the existing media as described in (35).

### Protein extraction, Western blot analysis and antibodies

Nuclear and cytoplasmic protein extracts were prepared as described in (37). Samples were analyzed by immunoblotting with mAbs targeted against p21 (N-20; Santa Cruz), PARP (9542; Cell Signalling), HuR (3A2; Santa Cruz), FLAG (4GFR; GeneTex), GAPDH (G9; Cell Signalling), Lamin A (H-102; Santa Cruz), CRT (F-4; Santa Cruz), CUGBP1 (B-1; Santa Cruz), GPD1 (HPA044620, Atlas Antibodies).

### 3’ Rapid Amplification of cDNA Ends (3’RACE)

Nuclear RNA was analyzed with the 3’ RACE System for Rapid Amplification of cDNA ends (ThermoFisher) as per the manufacturer’s protocol using oligo(dT)-adaptor primer (5’-GGCCACGCGTCGACTAGTACTTTTTTTTTTTTTTTTT-3’). PCR amplification was done using CDKN1A Exon 1 specific primer (5’-ATGCGTGTTCGCG GGTGT-3’) located 400 bp upstream of poly(A) site and adaptor (5’-GGCCACGCGTCGA CTAGTAC 3’). For nested PCR, a second CDKN1A specific forward primer in SPUD alternative exon was used (5’-AGCCGGAGTGGAAGCAGA-3’) and the same adaptor.

### Protein translation assays

FLAG-tagged SPUD *in vitro* expression was done with Transcend Non-Radioactive Translation Detection System (Promega) as per the manufacturer’s instructions. Expression was analyzed by chemiluminescence following a reaction of Lysine-biotin bound streptavidin-HRP to the substrate. *In vivo* expression was analyzed by Western blot with anti-FLAG.

### RNA immunoprecipitation (RIP)

The IP of nuclear RNA-protein complexes was performed as described (38). After treatments, cells were cross-linked and nuclear extracts were prepared. Extracts were treated with DNase (Ambion), and the resulting material was IP’ed with monoclonal antibodies against HuR (3A2; Santa Cruz), CRT (F-4; Santa Cruz), CUGBP1 (B-1; Santa Cruz) or control rabbit IgG (Sigma). Protein-RNA complexes were treated with proteinase K and reversal of cross-linking. The RNA was extracted from the IPs with phenol-chloroform and analyzed by RT-qPCR assays.

### RNA pull-down (RPD)

T7-driven pET-42a(+) construct containing SPUD transcript was used for *in vitro* transcription using RNA Biotin Labelling Mix (Sigma) and T7 polymerase (Promega) following manufacturer’s instructions. Biotinylated RNA was incubated with either nuclear extracts from HCT116 cells or recombinant His-HuR, GST-CRT, or GST-CUGBP1, and then pulled-down with magnetic streptavidin beads (ThermoFisher) followed by immunoblotting analysis, as described (38).

### Flow cytometry

Treated HCT116 cells were fixed and treated with staining mixture (0.1% Triton X-100, 2 mg RNAse A, 20 μg/ml propidium iodide). Samples were analyzed on FACSCalibur Flow Cytometer (BD Biosciences).

### RNA-seq/Ribo-Seq analysis

Publicly available RNA-seq and Ribo-seq dataset was downloaded from gene expression omnibus (NCBI) under accession number GSE99745. Downloaded fastq files were aligned to hg38 human genome using STAR and visualized using IGV (Broad Institute). Proportions of splice junctions used per sample were analyzed using IGV’s integrated sashimi plot function. Polysome to RNA-seq ratios were calculated by taking the junction depth of each polysome sample divided by the average junction depth across RNA-seq samples for three independent replicates (39,40). Footprint/RNA-seq ratios were calculated by the taking the value of peak coverage within the defined exon for each data track.

### Variant Analysis

Variants with CDKN1A intron 1 single nucleotide polymorphisms (SNPs) were searched using the genomic coordinates in the National Institutes of Health’s All of Us research database browser public tier(https://databrowser.researchallofus.org/). Detected variants within relevant sequences were visualized using Variation Viewer (NCBI).

## Results

### SPUD is a UV-inducible intragenic transcript generated by an intron-APA event in *CDKN1A* gene

Previously, we described a strong activation of intron-APA sites after UV-treatment, resulting in widespread expression of truncated transcripts, using 3’ region and extraction deep sequencing (3’READS, 32,41). Interestingly, these intron-APA events are biased to the 5’ end of genes and affect genes with functions in DDR and cancer. *CDKN1A* was detected to undergo UV-induced intron-APA in colon carcinoma RKO cells (32). As the coding sequence for p21 begins downstream in exon 2, the UV-induced APA event generated an entirely distinct, truncated transcript that we named Selective Polyadenylation Upon DNA Damage (SPUD). We then analyzed UCSC genome browser (42–43) to identify whether any annotated transcripts corresponded to SPUD transcript. Interestingly, a two-exon expressed sequence tag (EST; ENST00000462537) was detected terminating adjacent to the PAS identified by 3’READS in SPUD (Figure 1A, Supplementary Figure S1A, 32). The SPUD transcript shares exon 1 of CDKN1A full-length mRNA.

**Figure 1:**
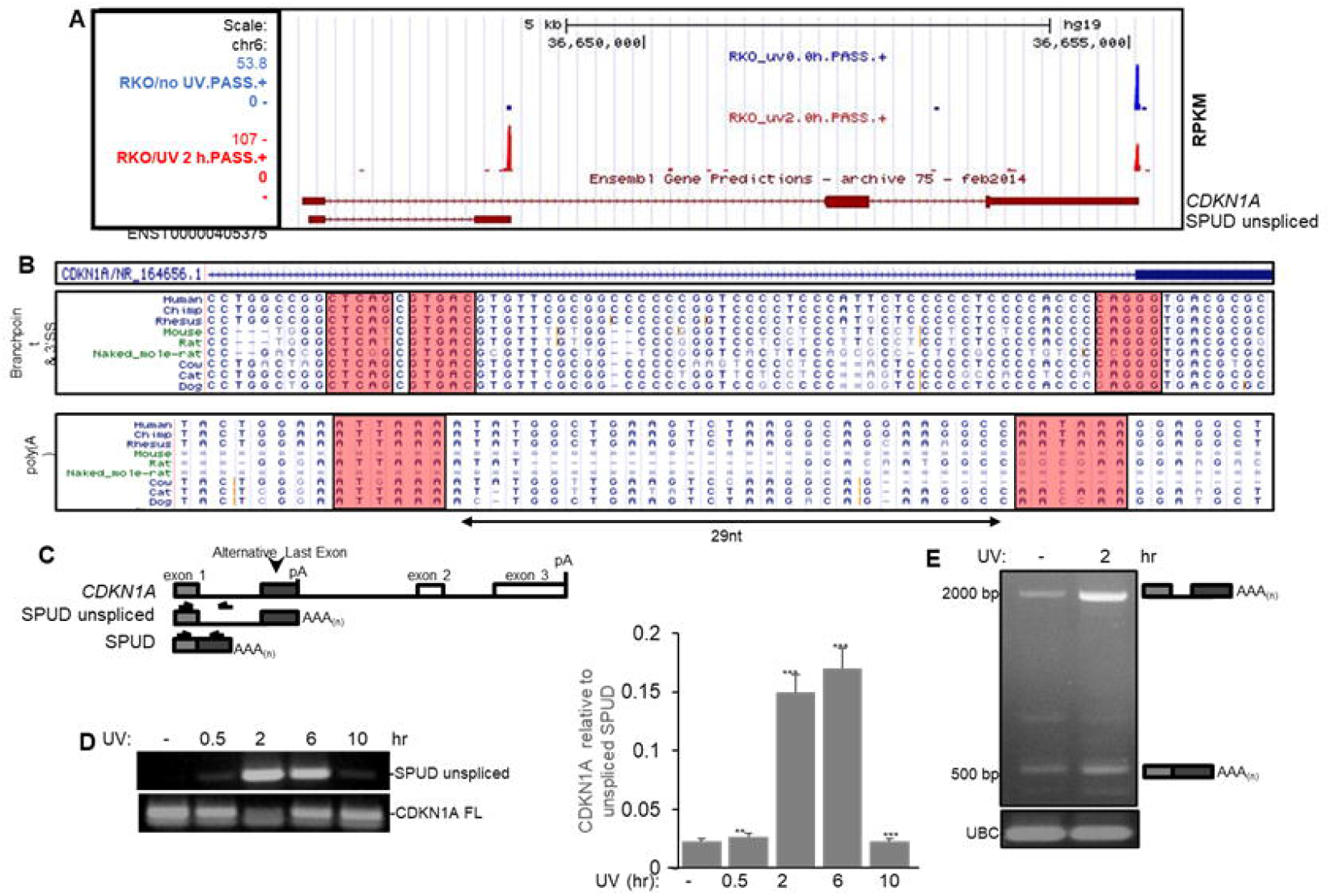
Transient UV-mediated APA event in CDKN1A intron 1 generates a processed transcript with an ALE. **(A)** Screenshot from the UCSC genome browser indicates an annotated transcript terminating at intron 1 (maroon diagram at the bottom). This intron-APA uses the same poly(A) signal in intron1 that the transcript detected via 3’ READS technique in samples from HCT116 cells treated with UV and allowed to recover for 2 h (20 J/m^2^). Detection was scored in Reads Per Million (RPM), calculated as the number of poly(A) site supporting (PASS) reads of hat site in a million unique PASS reads per sample. Blue and red bars indicate 3’READS CpA detection before and after UV treatment, respectively. Maroon boxes are exons and serrated lines are introns. Longer transcript on the top corresponds to CDKN1A full-length mRNA. **(B)** Multialignment of CDKN1A intron 1 in various mammalian species. Location and conservation of potential branchpoint and 3’ splice sites (3’ss) within CDKN1A intron 1 using screenshot from UCSC genome browser. CDKN1A intron 1 possesses two canonical PAS in proximity with one another with different evolutionary ages. Bar above represents the transcript for which the comparative analysis is examining. Highlighted light pink areas indicates the PAS and 3’ss aligned for each species. **(C)** Schematic of PCR strategies to detect CDKN1A full-length mRNA (top), unspliced intron-APA transcript (unspliced SPUD) and spliced intron-APA transcript (SPUD) for semi-quantitative and quantitative PCR. Isoform distinction is based on molecular weight. **(D)** Unspliced SPUD levels transiently increase after DNA damage without concomitant increase in full-length CDKN1A mRNA. Left: HCT116 cells were treated with UV (20 J/m^2^) and allowed to recover for indicated time points followed by RT-PCR using purified RNA and primer sets described in (B). cDNA was prepared using oligo(dT) primers. Right: ratio of unspliced SPUD to CDKN1A full-length mRNA. Values were normalized to non-treated cells. A representative gel from 3 independent biological samples analyzed by triplicate is shown. Errors represent SD (n=3). \*\**P*<0.001 and \*\*\**P*<0.0001. **(E)** Spliced transcripts generated from APA-ALE are detectable after UV treatment (20 J/m^2^, 2 h recovery) in samples from HCT116 cells by RT-PCR using SPUD primer set as in (B). Schematic next to gel indicates the SPUD isoform detected. Ubiquitin C (UBC) primers were used as loading control. A representative gel from 3 independent biological samples analyzed by triplicate is shown.

Further, inspection of ENST00000462537 indicated the presence of a canonical 3’ splice site (3’ss, CAG/GG, 44–45), which could be used to generate an ALE in SPUD, and the presence of two tandem PAS; AUUAAA and AAUAAA located ~57 nt and ~28 nt upstream, respectively, of the EST 3’ end (Supplementary Figure S1A, 26,46). Phylogenetic analysis across mammals for *CDKN1A* intron 1 PAS indicated that the upstream AUUAAA was conserved between rat and human, while the downstream AAUAAA was restricted to the simian lineage following an apparent C>T substitution (Figure 1B, Supplementary Figure S1C). Both signals were not present in every mammal tested, and the AUUAAA was absent in several species. The 3’ distal full-length mRNA PAS was almost invariably present in all mammals, except for a minority that possessed AGUAAA (Supplementary Figure S1C). A potential consensus branchpoint (44–45) was identified <60nt upstream of the 3’ss (Supplementary Figure S1B). Intriguingly, mutation of the critical nucleotides A or T, which are under strong evolutionary constraint for branchpoint formation (45), invariably was associated with alteration of the CAG/GG sequence of the 3’ss (eg. Marmoset).

SPUD isoforms, unspliced and spliced, were detected biochemically by RT-PCR using a forward primer targeting exon 1 and a reverse primer either in the intron or in the ALE (Figure 1C). Samples from HCT116 cells treated with UV irradiation (20 J/m^2^) and allowed to recover for the indicated time points were analyzed by RT-PCR. The levels of unspliced SPUD increased from 0 to 6 hours, reaching basal level detection by 10 hours (Figure 1D). The time-course for unspliced SPUD induction after DNA damage is similar to the time course previously described for cleavage/polyadenylation inhibition during DDR (47). Using the ALE reverse primer, RT-PCR analysis showed two bands that were increased by UV treatment (2 h recovery time, Figure 1E): a smaller band (~500 bp), which corresponded to SPUD (predicted spliced two-exon transcript; ENST00000462537), and a larger band (~2 kbp) that corresponded to unspliced SPUD or precursor. A third band was also detected (~750bp) that did not respond to UV treatment.

While both unspliced SPUD (~20 times) and SPUD (~8 times) were induced at early time points after UV treatment (2 h recovery), unspliced SPUD levels were twice of that observed for SPUD (Figure 2A). However, at later times post-UV (10 h recovery) SPUD levels almost doubled while unspliced SPUD reached basal levels, indicating that SPUD expression is sustained subsequent to transient APA. It is possible that unspliced SPUD is a truncated transcript produced by the previously described UV-induced block of transcription elongation somewhere in intron 1 (19,20,21,22), and that SPUD is the highly stable spliced product. Using the absolute Ct values from these experiments, we determined that CDKN1A full-length levels were ~120 times higher than SPUD prior to UV, and then reduced to ~16 times after UV treatment (Figure 2B). Taken together, this data describes for the first time an intron-APA event in the *CDKN1A* gene following DNA damage that generates a transcript with an ALE.

**Figure 2:**
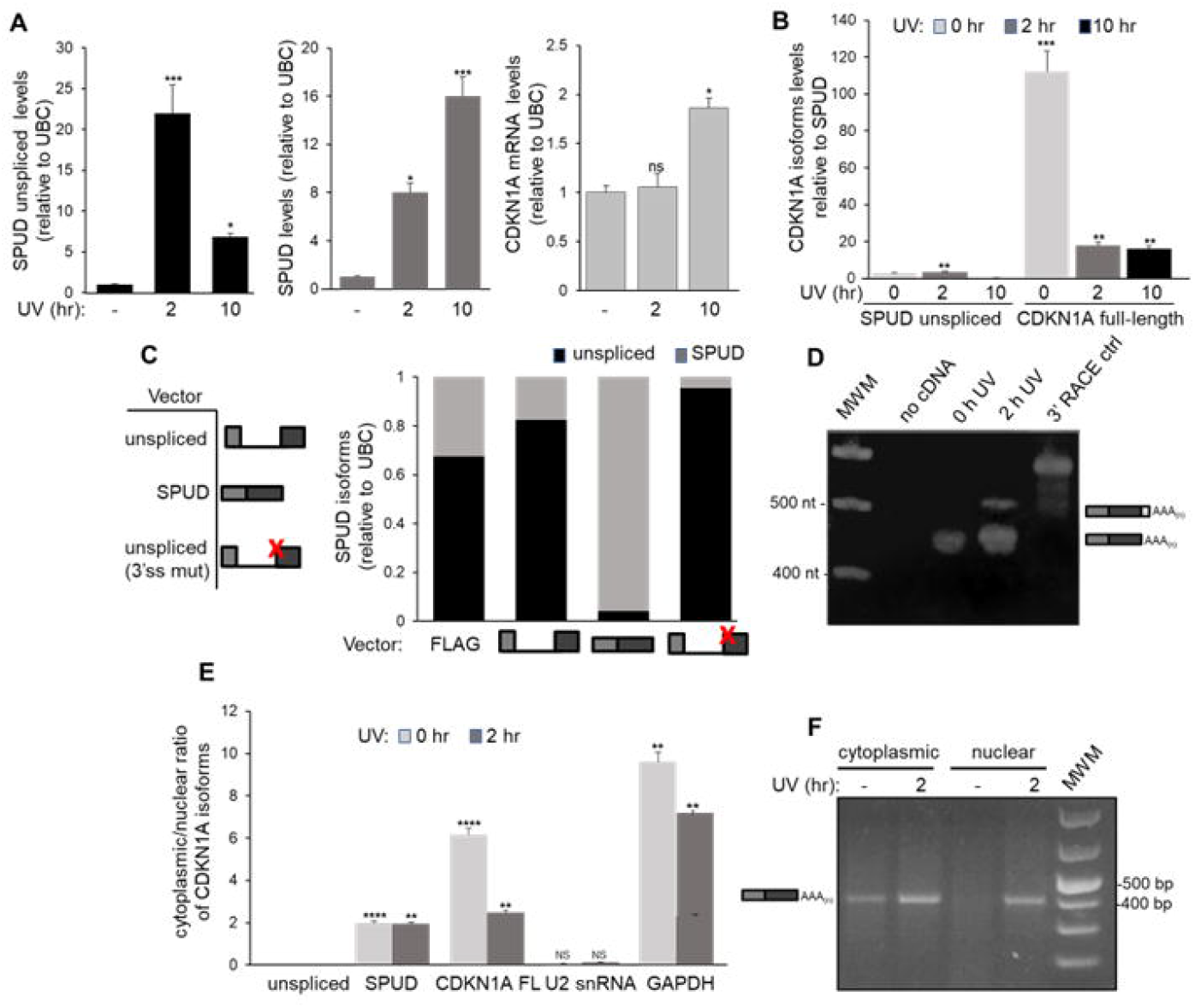
SPUD isoforms with multiple PAS and splicing elements and CDKN1A full-length expression levels show different patterns after UV-treatment. **(A)** While UV-transiently induces unspliced SPUD, SPUD can be detected longer in DDR progression. HCT116 cells were treated with UV (20 J/m^2^) and then recovered at times indicated followed by qRT-PCR using primers described in Supplementary Figure S5. cDNA was synthesized using oligo(dT) primers. The levels of the isoforms observed for each condition were normalized to the levels observed in non-treated cells. Three independent biological samples analyzed by triplicate is shown, SD (n=3). \**P*<0.01 and \*\*\**P*<0.0001. **(B)** CDKN1A full-length mRNA levels are higher than SPUD levels prior to UV, but then the molar difference between full-length mRNA and SPUD decreases after UV. SPUD levels are similar to unspliced SPUD in all the conditions. Ratios were calculated using the absolute Ct values from these experiments shown in (A) for the time points indicated. The ratios calculated for each condition were normalized to the ratios calculated in non-treated cells. \*\**P*<0.001 and \*\*\**P*<0.0001. **(C)** The predicted 3’ss is responsible for the detection of SPUD. Amplified cDNAs from UV-treated HCT116 cells were used to clone unspliced SPUD and SPUD into FLAG-tagged mammalian expression vectors. Unspliced SPUD vector was mutated to delete the CAGGG sequence shown in Figure 1B. Left: schematic of expression constructs; red ‘x’ indicating the vector with mutated 3’ss. Right: relative abundance of unspliced and SPUD in samples of each transfection condition was determined by qRT-PCR. cDNA was synthesized using oligo(dT) primers. Data shown is from 3 independent biological samples analyzed by triplicate, (n=3). **(D)** Both CDKN1A intronic PAS are used after UV treatment. Samples from UV-treated HCT116 cells were analyzed for CDKN1A intronic polyadenylation by 3’ RACE using a forward primer within exon 1 and a reverse oligo(dT)-tagged adapter primer. RNA was loaded as control for genomic DNA contamination. 3’RACE control cDNA provided by the kit was included. A representative gel from three independent assays from three biological samples is shown (n=3). **(E)** While unspliced SPUD localizes in the nucleus, SPUD localizes to the cytoplasm, and their distribution are unaffected by UV treatment. Nuclear and cytoplasmic RNA extraction was performed on samples from cells treated as in (A). qRT-PCR was performed as in (A). Cytoplasmic/nuclear ratio of 2^-ΔΔCT^ of equal RNA input. U2 snRNA and GAPDH were used as fractionation controls. Data shown is from 3 independent biological samples analyzed by triplicate, (n=3). \*\**P*<0.001 and \*\*\*\**P*<0.00001. **(F)** Spliced and polyadenylated SPUD is detectable in the cytoplasm after UV treatment. RNA extraction from cellular fractions was done as in (E) and 3’RACE analysis was done as in (C), except for an additional nested PCR using a downstream forward primer also within exon 1 for the second PCR. Samples were analyzed on an agarose gel and detected by ethidium bromide staining. A schematic of the product detected is shown adjacent. A representative gel from 3 independent biological samples analyzed by triplicate is shown. Molecular weight standard (MWS) is also included.

### SPUD is detected in the cytoplasm after undergoing splicing and polyadenylation

Furthermore, we characterized the RNA processing potential of SPUD. To determine 3’ss functionality, mammalian expression vectors were constructed for either the unspliced SPUD sequence, SPUD, or the unspliced sequence with a 5 nt deletion at the CAGGG splice site (Figure 2C). Samples from HCT116 cells transfected with the empty vector showed 2 fold difference unspliced SPUD/SPUD ratio (Figure 2C, first bar). As expected, overexpression of SPUD decreased that ratio by a factor of ten. Interestingly, overexpressing unspliced SPUD did not result in overexpression of spliced SPUD (4 fold difference), suggesting that the exogenously transcribed sequence was not efficiently spliced or that transcription outpaced the splicing rate. Unspliced SPUD/SPUD ratio was further increased with the expression of the splice-site mutant (10 fold), confirming the role of the intronic 3’ss in *CDKN1A* to generate the ALE in SPUD.

Next, we determined through 3’RACE which of the two tandem PAS was preferred in the mature SPUD transcript: the AUUAAA or AAUAAA located ~57 nt and ~28 nt upstream, respectively, of the EST 3’ end of SPUD. After UV treatment, we detected two bands; a stronger one consistent with the usage of the upstream AUUAAA PAS, and a band consistent with the usage of the downstream AAUAAA PAS Figure 2D), suggesting that the first PAS is preferred for the formation of the cleavage/polyadenylation complex. Interestingly, we identified human variants in unspliced SPUD-relevant regions, such exon 3’ss and PAS, using dbSNP repository and All of Us public tier websites from the NIH (Figure 3E). We found potentially inactivating rare variants for the 3’ss and both PAS of SPUD. While further experiments are necessary to test the effect of these variants on homeostasis, it is possible these non-coding variants have clinical significance.

**Figure 3:**
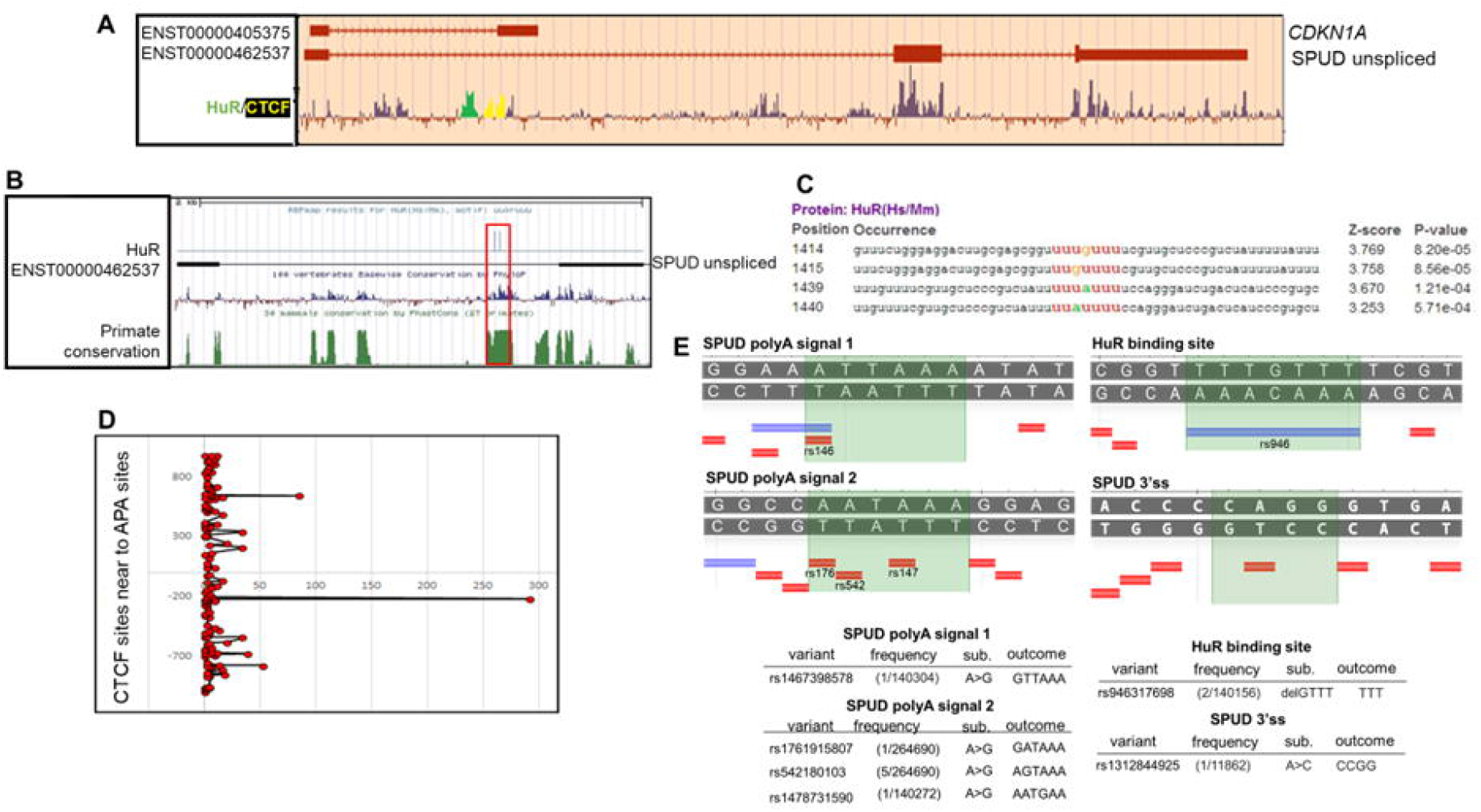
Predicted HuR and CTCF binding sites overlap with dense conserved regions in CDKN1A intron 1. **(A)** Screenshot of UCSC genome browser with multi-alignment of 100 vertebrate species showing regions of conservation along CDKN1A intron 1. Positive conservation as green peaks indicate region of conservation overlapping predicted HuR binding site. Positive conservation as yellow peaks indicate region of conservation overlapping CTCF binding site(s). Both sites are present within intron 1 overlapping EST corresponding to SPUD. **(B)** RBPmap motif analysis highlighting the two potential HuR binding sites corresponding to the region of conservation in (A). **(C)** RBPmap motif analysis identifies two AU-rich regions within CDKN1A intron 1 in close proximity with statistical likelihood to bind HuR. The entire 2 kbp sequence of unspliced SPUD event was analyzed with RBPmap for positions that overlapped with high vertebrate conservation as in (A). **(D)** Transcription factor CTCF binds to a sequence in region proximal to CDKN1A intron-APA site. Bioinformatic analysis of the distance of potential CTCF binding sites to PAS of genes undergoing intronic-APA. Each red dot indicates a different gene undergoing APA, while its corresponding black line represents the expression levels of that APA isoform after UV treatment. (*) indicate CDKN1A intron-APA. **(E)** Variants with CDKN1A intron 1 SNPs were searched using the genomic coordinates in the NIH’s All of Us research database browser public tier and dbSNP repository and were visualized using Variation Viewer (NCBI).

As splicing and polyadenylation have been previously described to stimulate nuclear export (48), we next tested SPUD localization. As expected, qRT-PCR analysis of subcellular RNA preparations showed U2 snRNA and GAPDH mRNAs to be predominantly nuclear and cytoplasmic, respectively (Figure 2E). While CDKN1A full-length mRNA was 6-fold enriched in the cytoplasm, this enrichment decreased upon UV treatment. Unspliced SPUD was exclusively nuclear, as expected for an unspliced RNAs (49). SPUD was enriched in the cytoplasm but at lower levels than the full-length mRNA (Figure 2E). While SPUD abundance increased after UV treatment, cytoplasmic/nuclear ratio was unchanged. Consistent with this, 3’RACE with nested PCR showed that SPUD was predominantly cytoplasmic and was induced after UV treatment in both cytoplasmic and nuclear fractions (Figure 2F). Together, these results indicate that SPUD is a cytoplasmic RNA that is subject to a different regulatory processing than the coding CDKN1A mRNA under stress conditions.

### RNA binding protein (RBP) HuR and the transcriptional repressor CCTC-binding factor (CTCF) regulate SPUD levels

Previously, it was described that inhibition of the splicing regulator U1 snRNA function leads to activation of intron-APA events, resulting in shorter transcripts (30–31), and that U1 RNA levels transiently decrease upon UV treatment (50). Work from our lab showed that U1 was also implicated in the CDKN1A intron-APA event as functional depletion of U1 RNA using morpholino oligonucleotides (AMO) increases the SPUD/CDKN1A full-length ratio. Interestingly, the changes in the SPUD/CDKN1A full-length ratio by U1 RNA depletion were similar in magnitude to that observed after UV treatment. However, there is a discrepancy in the timely and controlled regulation of these processes. While U1 snRNA levels decline between 2-6 h post-UV, the levels of SPUD remain constant (32), suggesting that gene-specific regulators of SPUD might play a role. Analysis of vertebrate conservation within the unspliced SPUD transcript indicated two regions of positive selection <500 bp upstream from the SPUD 3’ss, a binding region for the transcriptional repressor CTCF and for the RBP HuR (Figure 3A, Supplementary Figure S2A). Both pre-existing ChIP-seq data from ENCODE (51) and a DNase I hypersensitive site across multiple cell lines (Supplementary Figure S2B) also detected CTCF binding in the 3’ss proximal region of unspliced SPUD. RBPmap, a program that predicts the likelihood of RBPs interacting with a subject RNA (52–53), detected two motifs for HuR within the ~2 kbp unspliced SPUD (Figure 3B) using the consensus sequence “uukruuu” (Figure 3C), which has been identified as a U-rich sequence required for HuR binding (54). Interestingly, we identified a deletion variant located at the HuR binding site (Figure 3E). Further experiments are necessary to test whether this non-coding variant has clinical significance.

Recent research uncovered a link between CTCF binding and APA events by binding downstream of promoter-proximal PAS (55). Bioinformatic analysis of intron-APA sites relative to CTCF sites showed that the CDKN1A intron-APA site located ~200 bp downstream of a CTCF site had the strongest induction of all APA sites in proximity to CTCF (Figure 3D). Interestingly, CTCF depletion upon UV treatment (Supplementary Figure S2C) increased the levels of both SPUD and CDKN1A full-length mRNA but not of unspliced SPUD (Figure 4A). The increase in CDKN1A fulllength mRNA is consistent with CTCF role in transcription repression (56). However, as SPUD and unspliced SPUD originate from the same transcriptional unit, these results suggest that CTCF might play a role in promoting inclusion of a weak ALE in SPUD during stress conditions. The lack of unspliced induction, despite SPUD upregulation, suggests rapid splicing upon CTCF knockdown. Supporting our results, it has been shown that CTCF can induce inclusion of weak upstream exons by mediating local RNA polymerase II pausing and allow co-transcriptional RNA processing both in a mammalian model system for alternative splicing and genome-wide (57).

**Figure 4:**
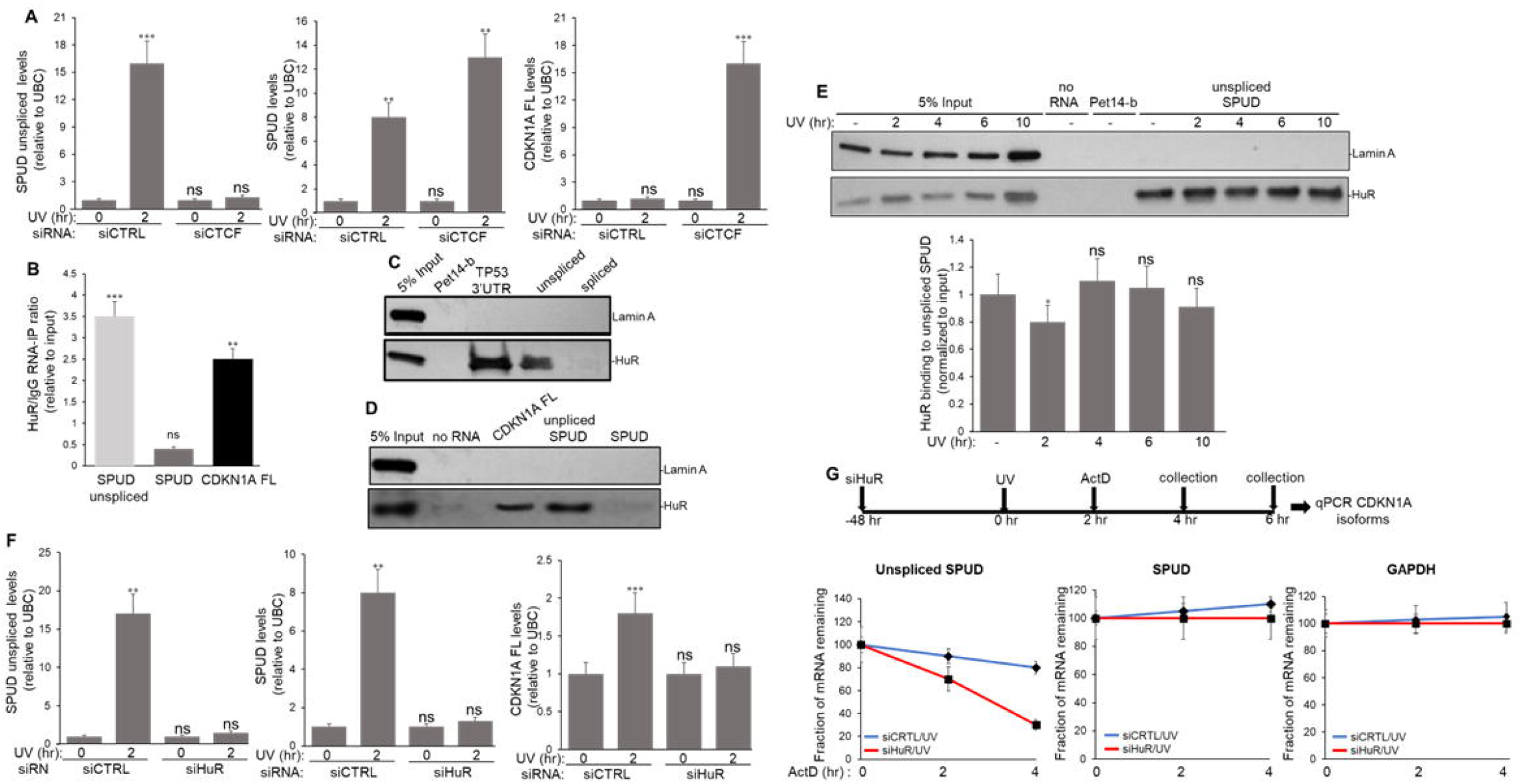
While HuR interacts and regulates the stability of unspliced SPUD independently of DNA damage, CTCF downregulates SPUD but not the unspliced isoform. **(A)** siRNA-mediated depletion of CTCF results in upregulation of SPUD and CDKN1A full-length, but not the unspliced isoform. HCT116 cells were treated with control (CTR) or CTCF siRNA and then with UV (20 J/m^2^), allowed to recover for 2 h followed by RNA extraction. CDKN1A isoforms were analyzed by qRT-PCR as in Figure 2A. The levels of the isoforms observed for each condition were normalized to the levels observed in non-treated cells. Data shown is from 3 independent biological samples analyzed by triplicate, SD (n=3).\*\**P*<0.001 and \*\*\**P*<0.0001. **(B)** HuR binds to unspliced but not SPUD isoform. HCT116 cells were treated with formaldehyde to generate protein-RNA cross-links. The samples were incubated with either anti-HuR or IgG antibodies. The endogenous nuclear RNA IP’ed with the antibodies was quantified by qRT-PCR using primers specific for each CDKN1A isomers described in Figure 1C. The qRT-PCR values were calculated from 3 independent biological samples analyzed by triplicate, (n=3). The input indicates samples before RIP. \*\**P*<0.001 and \*\*\**P*<0.0001. (**C-D**) Unspliced but not spliced SPUD interacts with HuR independently of UV treatment. RNA pull-down assays were performed using *in vitro* transcribed biotinylated RNA from the CDKN1A APA isoforms mixed with NEs from HCT116 cells. Equivalent amounts of the eluate were resolved by SDS-PAGE and proteins were detected by immunoblotting using antibodies against Lamin A and HuR. *In vitro* transcribed biotinylated Pet14-b RNA was used as nonspecific sequence (no HuR sequence is present in this fragment), while RNA from TP53 and CDKN1A 3’UTRs were used as positive control for binding (95). Lamin A was used as negative control for binding. A representative gel from 3 independent biological samples analyzed by triplicate is shown. Five percent of the NEs used in the pull-down reactions is shown as input. (**E**) HuR binding to unspliced SPUD is independent of UV treatment. Top: NEs from HCT116 cells exposed to UV (20 J/m^2^), allowed to recover for the indicated times were incubated with biotinylated SPUD and proteins were detected as in (B). Bottom: quantification of RNA pull-down assays represented in the top. The levels of HuR pulled down in each condition were normalized to the levels observed in each input. A representative pull-down reaction from 3 independent biological samples analyzed by triplicate is shown. Five percent of the NEs used in the pull-down reactions is shown as input. \*\**P*<0.001 (**F**) siRNA-mediated knockdown of HuR prevents UV-induced activation of CDKN1A isoforms. RNA levels of SPUD isoforms and fulllength CDKN1A mRNA were analyzed by qRT-PCR in samples from cells treated with HuR/control siRNA for 48 h and UV irradiation (20 J/m^2^, 2 h recovery). RNA abundances were normalized to UBC. Data shown is from 3 independent biological samples analyzed by triplicate, SD (n=3). \*\**P*<0.001 and \*\*\**P*<0.0001 (**G**) HuR regulates abundance of CDKN1A APA isoforms in transcription-independent manner. Unspliced SPUD stability is reduced upon HuR depletion. HCT116 cells treated with HuR/control siRNAs and UV irradiation as in (F). Cells were also incubated with actinomycin D for the indicated times after UV treatment followed by qRT-PCR of SPUD isoforms. mRNA decay rates for SPUD isoforms and GAPDH, a non-HuR target transcript, were determined by qRT-PCR at different time points following HuR/control siRNA-, UV- and Act-D treatment. The relative half-life of the SPUD isoforms transcript was calculated from three independent samples. Errors represent the SD derived from three independent experiments.

HuR can bind not only the 3’ untranslated region (3’UTR) of full-length mRNAs, regulating their stability (35,58), but also lncRNAs, affecting their stability and localization (59–60). In fact, intronic HuR sites are more prevalent than 3’UTR binding regions (61). RNA-immunoprecipitation (RIP) assays showed that HuR can form complexes with both CDKN1A full-length mRNA and unspliced SPUD but not with SPUD (Figure 4B). The intronic sequence in unspliced SPUD was required to form a complex with HuR from HCT116 nuclear extracts (NEs) in RNA-pull down (RPD) assays with biotinylated CDKN1A isoforms (Figure 4C-D). When NEs from cells recovered at different time points after UV treatment were used in RPD assays, HuR binding to unspliced SPUD was largely unchanged (Figure 4E), except for a small but significant decrease at early time points after UV treatment. Importantly, unspliced SPUD-HuR binding was detected at the 10 h time point, at a time when the unspliced SPUD levels started to decrease to non-stress levels (Figures 1D and 2A). However, SPUD levels were still high, suggesting that HuR binding to unspliced SPUD is likely not implicated in regulating the APA event itself as has been previously described for others (62). Interestingly, HuR depletion upon UV treatment (Supplementary Figure S2D) abolished not only the previously described increase in CDKN1A full-length mRNA (35) but also induction of both unspliced SPUD and SPUD (Figure 4F, Supplementary Figure S2E). As HuR is a highly abundant nuclear protein (63), overexpression of HuR did not affect the UV-induced increase in CDKN1A full-length mRNA, unspliced SPUD and SPUD (Supplementary Figure S2F).

To further understand the mechanism for this HuR-mediated regulation of CDKN1A isoforms, HCT116 cells were pre-treated with UV with a 2 h recovery to induce APA followed by actinomycin D treatment for the time course indicated (Figure 4G). Intriguingly, HuR depletion resulted in unspliced SPUD half-life decreased from 8.1 h to 2.8 h, while control GAPDH mRNA half-life did not change. Notably, a downward trend in SPUD was observed upon HuR depletion, but this was not significant. Overall, these results, indicate that HuR regulates the stability of unspliced SPUD, adding another lncRNA to HuR’s regulatory repertoire (13).

### SPUD is induced in cancer and normal cells under a variety of damaging conditions in a p53-dependent manner

To further understand whether SPUD induction was cell type- or stress-specific, we extended our studies to other cell lines and analyzed the effect of different stressors. A delay in *CDKN1A* mRNA and p21 protein expression due to a block in transcription elongation has been described under certain stressors, such as UV and HU treatment (19,20,21,22), but not others, such as the strong p21 inducer, etoposide (64). It is possible that SPUD induction might also occur in certain stress conditions, but not others, reflecting its cellular function in DDR. Interestingly, expression of both SPUD and full-length mRNA was induced upon etoposide treatment, suggesting that the role of SPUD is not to suppress *CDKN1A* full-length expression (Figure 5A-B).

**Figure 5:**
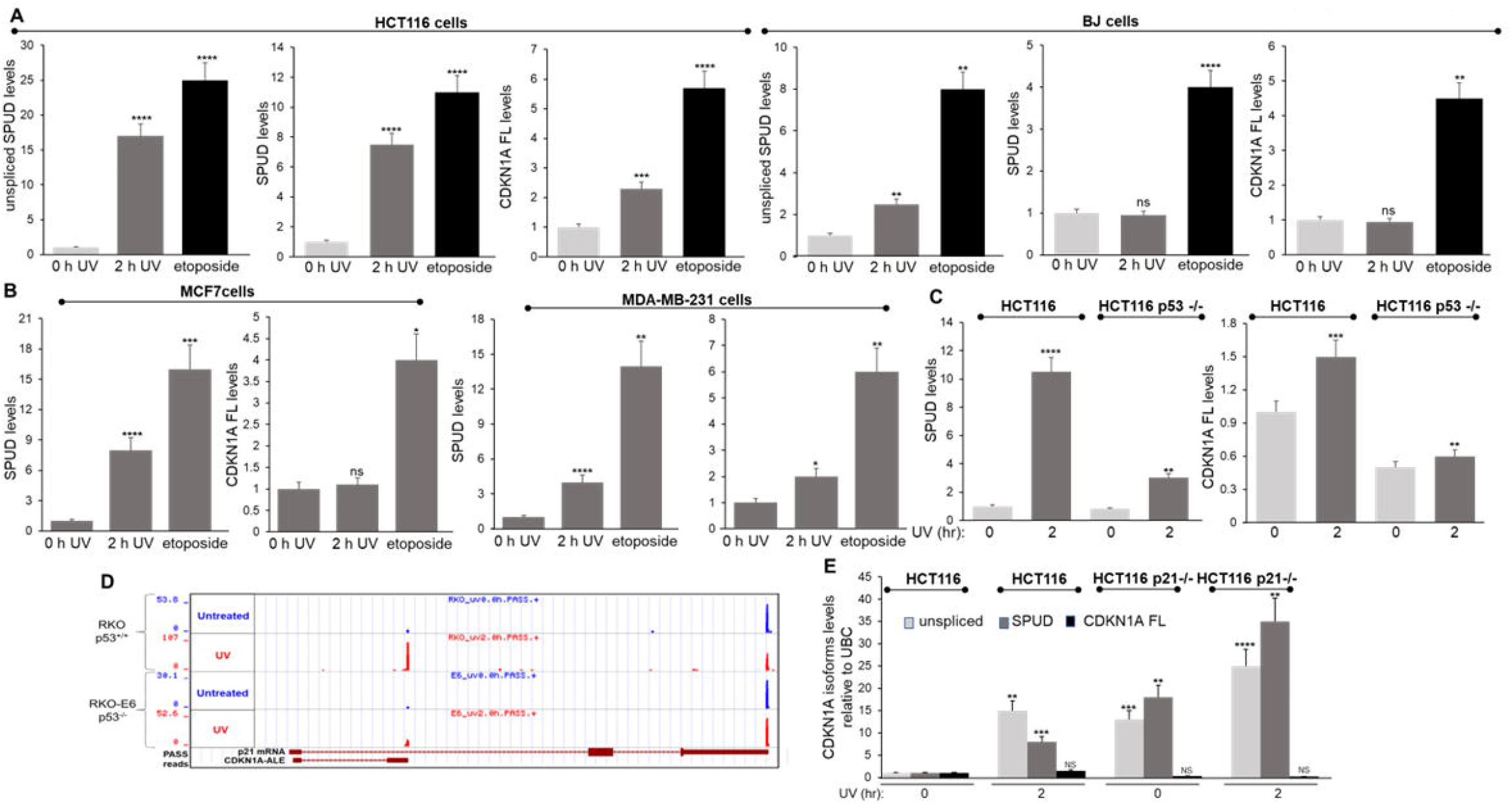
SPUD isoforms are expressed in a variety of cell lines and stressor conditions in a p53-dependent manner. (**A**) Induction of SPUD was higher in tumorigenic cell lines (HCT116) than into non-cancer cell lines (BJ-5ta immortalized foreskin fibroblasts). Total RNA was purified from the indicated cell lines and analyzed by qRT-PCR as in Figure 2A. Cells were treated with UV irradiation (20 J/m^2^, 2 h recovery) or with etoposide (20 μM) 16 h prior to total RNA extraction. Data shown is from 3 independent biological samples analyzed by triplicate, SD (n=3). ** p<0.005, *** p<0.0005, **** p<0.00005. (**B**) MCF7 or MDA-MB-231 breast cancer cells were treated and analyzed as in (A). (**C**) Loss of p53 significantly reduces expression of both SPUD isoforms and CDKN1A full-length. HCT116 and HCT116 p53^-/-^ cells were treated and analyzed as in (A). (**D**) 3’READS data indicates ablation of intron-APA in p53-null cells. Analysis of RKO and p53-null RKO-E6 colon carcinoma cell lines before (blue) and after (red) UV-mediated DNA damage. Detection was scored in Reads Per Million (RPM), calculated as the number of poly(A) site supporting (PASS) reads of hat site in a million unique PASS reads per sample. (**E**). Unspliced SPUD and SPUD expression is detectable and dysregulated in HCT116 p21^-/-^ cells. Both HCT116 and HCT116 p21^-/-^ cell lines were treated and analyzed as in (A).

SPUD generated by UV-induced intron-APA in CDKN1A was first described in colon carcinoma RKO cells (32). Here we also show that SPUD can be detected in HCT116 colorectal carcinoma (Figures 1–4), a cell line historically used to study *CDKN1A* transcriptional regulation (17,32). While SPUD was also inducible after UV and etoposide treatments in MCF7 breast cancer and MDA-MB-231 triple-negative breast cancer cell lines (Figure 5B, Supplementary Figure S3B), CDKN1A full-length was only induced in MDA-MB-231 upon UV treatment. To test whether SPUD is the result of aberrant APA in a tumorigenic background, we assessed UV-induced increase in SPUD in a noncancerous, physiologically relevant cell line, immortalized skin fibroblast cell line BJ1-hTERT. Interestingly, while UV-induction of SPUD was not detectable BJ1 at 2 h recovery time, activation of both SPUD and CDKN1A full-length occurred upon etoposide treatment in BJ1 cells (Figure 5A), suggesting that UV-mediated induction of SPUD is a widespread, common feature of DDR in many different cultured human cells with different tissues of origin.

Our previous 3’ READs data generated in RKO-E6 cells, which lack functional p53, indicated that many intron-APA events were directly or indirectly p53-dependent, including *CDKN1A* intron-APA (32). As SPUD is a sense-strand intragenic transcript of *CDKN1A* gene, we postulated that SPUD transcriptional induction might be subject to the same promoter regulation by p53 (65). Indeed, UV-induced expression of both SPUD and, as previously described (19,20,21,22,66), *CDKN1A* full-length decreased, but was not entirely abolished, in HCT116 p53 -/- cells (Figure 5C). This is consistent with our previous high-throughput screen that showed the reduction in PAS supporting (PASS) reads is partly dependent on p53 (Figure 5D, 32). Notably, a previously derived cell line lacking p21 expression due to recombination-mediated excision of coding exons 2 and 3 (67) exhibited significantly higher SPUD levels at baseline as well as augmented UV-mediated induction of SPUD but not of *CDKN1A* full-length (Figure 5E). Together, these results indicate that SPUD is expressed across a range of cell-types and treatments and likely depends upon the same p53 responsive elements as *CDKN1A* promoter.

### SPUD is a lncRNA that regulates the translation of p21 protein

An intriguing aspect of CDKN1A full-length mRNA is that the coding sequence begins in exon 2, downstream of the APA signal (Figure 6A), but that the mRNA also has an inhibitory upstream open reading frame (uORF) ATG in exon 1. PhyloCSF analysis of *CDKN1A* gene confirmed the positive selection for CDKN1A full-length mRNA ORF (Figure 6A, 70,71). Average PhyloCSF score for SPUD-ALE in the same frame as uORF ATG was −4.43, despite the presence of a 3’ss branchpoint and ATTAAA PAS in both rats and humans (Supplementary Figure S1B-C). To test experimentally whether SPUD coded for protein, expression was analyzed in samples of HCT116 cells transfected with mammalian vectors with N-terminal FLAG tag adjacent to unspliced precursor or SPUD cDNA encompassing the full-length uORF ATG. While FLAG immunoblotting detected a band for the FLAG-GPD1 control, it did not identify a specific band for SPUD isoforms (Figure 6B, left). Additionally, *in vitro* translation of SPUD constructs with biotinylated lysines did not yield detectable protein, while a band was detected for an HuR positive control construct (Figure 6B, right). Thus, these results are consistent with the idea that SPUD is a putative intragenic lncRNA within the *CDKN1A* gene.

**Figure 6:**
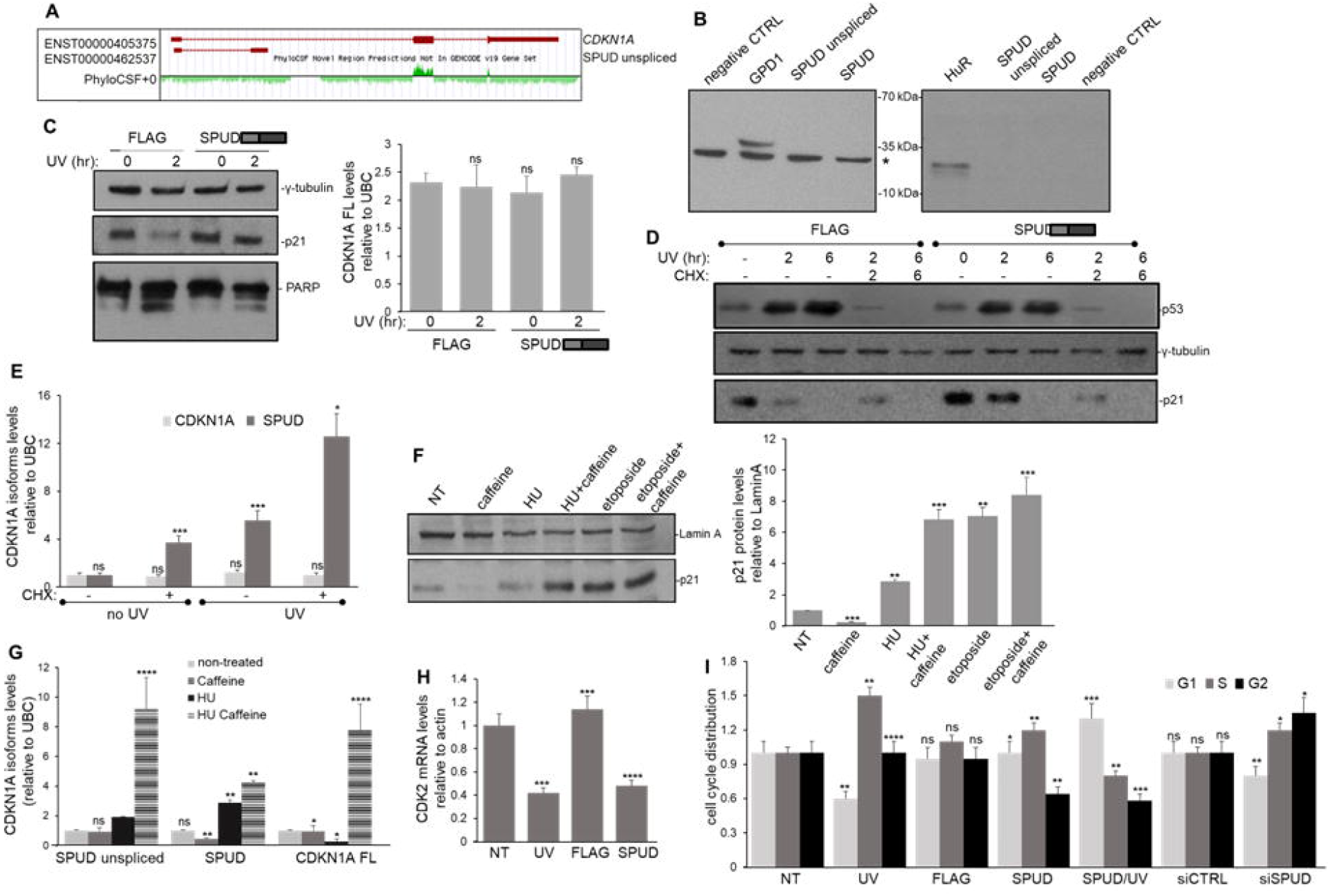
SPUD is a lncRNA that regulates p21 post-transcriptionally affecting p21 functions in cell-cycle, CDK2 expression, and cell viability. (**A**) PhyloCSF shows negative average score for CDKN1A exon 1 and intron 1, whereas p21 exons 2 and 3 have positive score. UCSC genome browser screenshot of PhyloCSF analysis of CDKN1A. Positive green peaks indicate coding sequence conservation, negative peaks indicate no positive selection for coding sequence. (**B**) *Ex vivo* and *in vitro* translation reactions yield no detectable protein product for both unspliced SPUD and SPUD isoforms. Left: 0verexpression of SPUD isoforms make no detectable protein in HCT116 cells. CMV-driven expression plasmids with N-terminal FLAG tags containing no transcript, GDP1 as positive control, or either unspliced SPUD/SPUD transcripts were overexpressed. Total protein was extracted from transfected cells and immunoassayed with anti-FLAG antibodies. (*) indicates non-specific band detected by the antibody. A representative gel from three independent samples is shown. Right: *In vitro* translation reactions yield no detectable protein for SPUD isoforms. T7-driven plasmids containing HuR as a positive control or unspliced SPUD/SPUD transcripts were *in vitro* translated in rabbit reticulocyte lysates. Translated proteins were labelled with biotinylated lysines, reactions were run on SDS-PAGE and immunoassayed against streptavidin-HRP conjugates. A representative gel from three independent samples is shown. (**C**) SPUD overexpression upregulates p21 and mitigates UV-mediated transient downregulation of p21 protein without affecting CDKN1A full-length mRNA levels. Cleaved PARP is detected as a biomarker for apoptosis. Left: HCT116 cells were transfected with CMV-driven expression plasmids with FLAG-tagged SPUD transcript and exposed to UV irradiation (20 J/m^2^, 2 h recovery). NEs were immunoassayed for p21, PARP and γ-tubulin as control. Gel shown is representative of three independent biological samples. Right: RNA from transfected HCT116 cells with expression constructs described above was analyzed by qRT-PCR for CDKN1A full-length mRNA and normalized to UBC. Data shown is from 3 independent biological samples analyzed by triplicate, SD (n=3). ns: no significant. (**D**) Inhibiting translation with CHX prevents the effect of overexpressed SPUD on the reduction in UV-induced decrease in p21, but has no effect on p53 levels. HCT116 cells were transfected as in (C) and treated with either UV (20 J/m^2^) or CHX (2 μg/ml)/UV (20 J/m^2^) for the indicated times. Samples were analyzed as in (C). (**E**) Inhibiting translation with CHX leads to higher SPUD induction after UV treatment. Cells were treated as in (D) for RNA purification analyzed by qRT-PCR for CDKN1A isoforms as in Figure 2A. (**F**) Caffeine treatment rescues p21 induction following S-phase block with HU. Left: HCT116 cells were treated with HU (1.7 mM) or etoposide (20 μM) 16 h followed by 2 h incubation with caffeine (4 mM) prior to protein (F) or RNA (G) purification. F-Left: protein samples were immunoassayed for p21 and Lamin A as control. Gel shown is representative of three independent biological samples. F-Right: quantification of protein expression shown in the left. Data shown is from 3 independent biological samples analyzed by triplicate, SD (n=3).\*\**P*<0.001 and \*\*\**P*<0.0001. (**G**) Rescuing full-length CDKN1A mRNA induction with caffeine treatment does not abrogate DNA-damage induced SPUD. RNA samples were were analyzed by qRT-PCR for CDKN1A isoforms as in Figure 2A. (**H**) Overexpression of SPUD leads to CDK2 mRNA downregulation. HCT116 cells were treated as in (C) and analyzed for CDK2 transcript by qRT-PCR as in Figure 2A. (**I**) Altering SPUD expression affects cell cycle distribution. HCT116 cells were treated for SPUD overexpression as in (C) or for SPUD depletion by siRNA - mediated knockdown as in Figure 7. Cells were fixed with 4% formaldehyde, permeabilized with 95% ethanol and then treated with propidium iodide (PI) for FACS analysis. Data shown is from 3 independent biological samples analyzed by triplicate, (n=3). ns; not significant; \**P*<0.01 \*\**P*<0.001; \*\*\**P*<0.0001 and \*\*\*\**P*<0.00001.

Several notable examples of intragenic/intronic lncRNAs have been described working either *in cis* or *in trans* to affect the expression of the protein of their host protein-coding gene (5,72). Therefore, we examined the role of SPUD on p21 protein expression levels, knowing that a downregulation occurs at early time points after UV treatment (Figures 6C-D, 4). We found that samples from cells overexpressing SPUD mitigated UV-induced downregulation of p21 expression (Figure 6C, left), and this was independent of CDKN1A full-length mRNA levels (Figure 6C, right). An upregulation of p21 was also observed when expressing the unspliced plasmid (Figure S3C). In addition, while UV-induction increased levels of cleaved PARP, which was shown to be inhibited by p21 expression (73), SPUD upregulation in the presence of UV diminished PARP cleavage (Figure 6C). These results suggest that SPUD works *in trans* as a p21 translational regulator, which is also supported by its cytoplasmic localization (Figure 2E). Consistent with this, treatment of cells with cycloheximide, a translational inhibitor, prevented the enhanced expression of p21 that occurred during exogenous induction of SPUD (Figure 6D). SPUD did not exert a regulatory feedback loop over p53 expression, as overexpression of SPUD did not alter p53 levels (Figure 6D). Notably, cycloheximide treatment combined with UV led to even greater induction of SPUD (Figure 6E), indicating that SPUD induction but not p21 expression is independent of translational activity.

Prior research has shown that CDKN1A full-length transcription is inhibited at the level of elongation by certain treatments, such as UV and HU, but not topoisomerase II inhibitors, such as etoposide (19,22). Transcription elongation block occurs in part by DDR signaling kinase Chk1 (22), and this can be inhibited by caffeine pretreatment resulting in the rescue of transcription elongation of CDKN1A full-length(22). We proposed that the release of transcription blockage would also increase the levels of SPUD isoforms. As previously shown (22), our results indicate that caffeine treatment in the absence of a stressor decreases the levels of p21 (Figure 6F). HU treatment, like UV irradiation, induced SPUD isoforms with a delay in full-length mRNA levels. Caffeine pretreatment in combination with HU induced not only protein expression (Figure 6F) but also the levels of all CDKN1A transcript isoforms (Figure 6G). Caffeine pretreatment did not significantly increase etoposide’s effect on p21 levels (Figure 6F). Therefore, rescuing transcription elongation of full-length mRNA also led to an upregulation of SPUD and its unspliced precursor. While further studies are needed, it appears that Chk1-mediated transcription elongation inhibition has an effect on SPUD activation, providing further support that CDKN1A and SPUD isoforms originate from the same transcriptional unit.

As previously described (74), flow cytometry analysis of UV-treated cells showed an increase in S-phase and a decrease in G1 due to p21 decrease (Figure 6I). Overexpression of SPUD under UV treatment led to an increase in G1 and decrease in S/G2 phase, as expected for induction of p21 protein expression. Likewise, expression of SPUD also led to a decrease in CDK2 mRNA (Figure 6H, 75). Together, these studies indicate that SPUD can induce functional p21 protein during DDR. Importantly, depletion of SPUD and unspliced SPUD using an siRNA targeting the APA exon (siSPUD, (Figure 7A-C, ~60% decrease) did not significantly change the levels of CDKN1A full-length mRNA (Figure 7B) but did decrease p21 protein levels in both cytoplasmic and nuclear fractions (Figure 7C). SPUD depletion led to a decrease in G1 and increase in S/G2 phase, as expected for a decrease in p21 protein expression (Figure 6I). Consistent with p21’s role as an inhibitor of cellular proliferation (76), SPUD depletion led not only to a decrease in p21 levels (Figure 7C) but also to an increase in the cell count compared to siCtrl treated cells (Figure 7D), indicating a loss in cell-cycle arrest capabilities. Together these results indicate that SPUD has a positive regulatory role on p21 translation.

**Figure 7:**
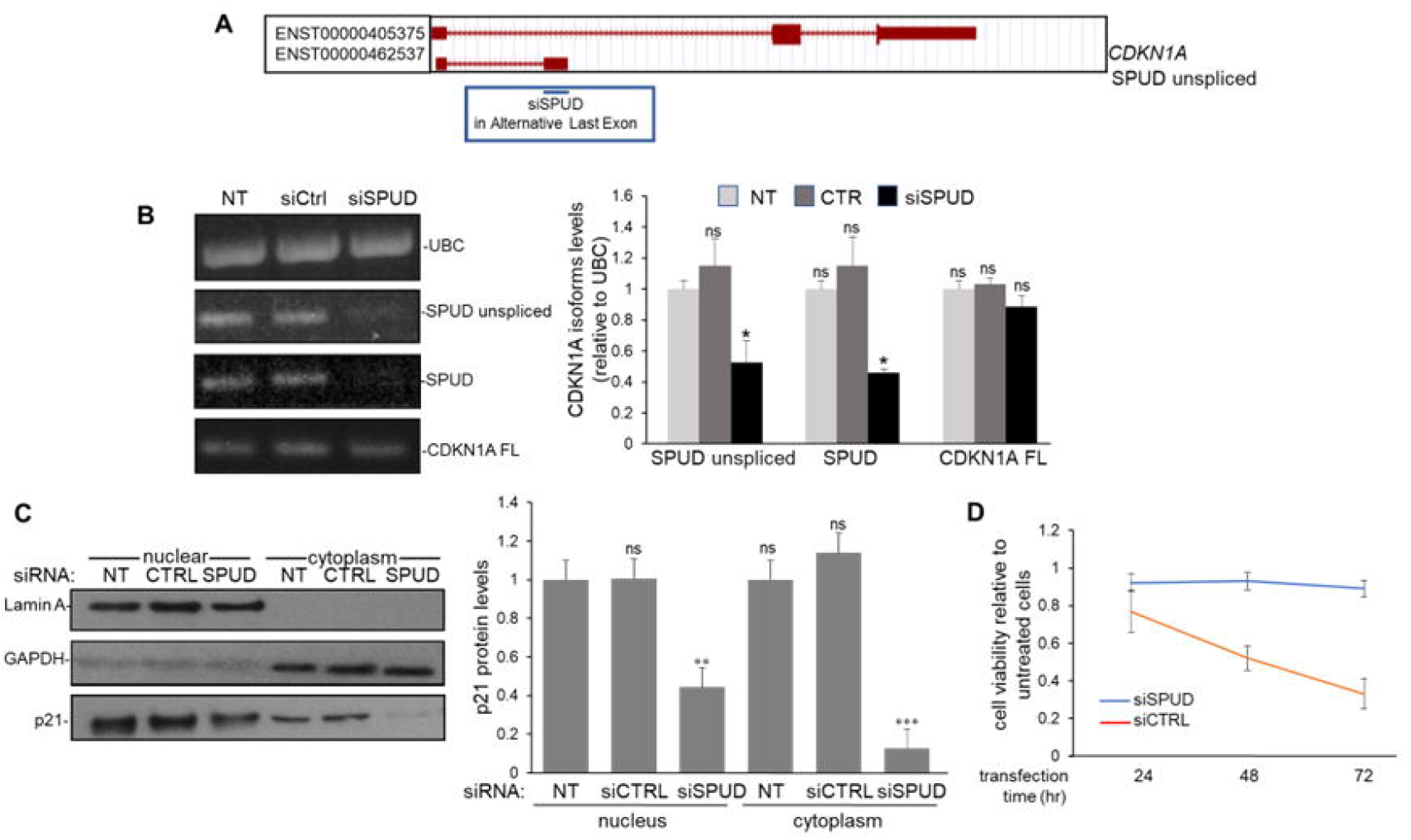
SPUD depletion decreases p21 protein expression without significantly affecting CDKN1A full-length mRNA levels. (**A**) Schematic of the design of an siRNA to target SPUD (siSPUD). Custom siSPUD was designed using siDESIGN Center (Horizon Discovery) targeting the entire APA exon as input. The black line represents the location of siSPUD in SPUD-ALE, the representation is not to scale. (**B**) siSPUD can deplete SPUD expression without significantly affecting CDKN1A full-length expression. RNA from HCT116 cells treated with siSPUD were analyzed by semi-quantitative PCR (left) or qRT-PCR (right). PCR products were separated on an agarose gel and detected by ethidium bromide staining. UBC amplification was used as loading control. Shown is a representative gel from three independent assays. qRT-PCR for CDKN1A isoforms was performed as in Figure 2A. Data shown is from 3 independent biological samples analyzed by triplicate, (n=3). ns; not significant; \**P*<0.01. (**C**) SPUD depletion reduces the total abundance of cellular p21 protein. Left: Nuclear and cytoplasmic protein fractions were prepared from cells treated as in (B). Samples were analyzed by immunoblotting for p21 expression. Lamin A and GAPDH were used as fraction control. NT, non-transfected; CTRL, control siRNA. Right: SPUD knockdown has greater impact on cytoplasmic than nuclear p21. Data shown is from 3 independent biological samples analyzed by triplicate, (n=3). ns; not significant; \*\**P*<0.001 and \*\*\**P*<0.0001. (**D**) SPUD depletion diminished loss of viability caused by control siRNA transfection. Cells were transfected as in (C) and measured via trypan blue at the indicated timepoints post-transfection. Data shown is from 3 independent biological samples analyzed by triplicate, (n=3).

### Different SPUD spliced isoforms associate with ribosome

Both ribosome profiling (77) and polysome profiling (78) studies have highlighted that approximately half of the annotated lncRNAs localize in the cytoplasm, most of which are associated with ribosomes. Analysis of a pre-existing RNA-seq dataset of UV-treated N-TERT keratinocytes showed a strong induction of SPUD after 8 h of UV-B treatment relative to full-length (100 J/m^2^, Figure 8A; 39). While the cell lines and stress conditions are different, these results are consistent with the strong induction beyond baseline detected for SPUD 10 h post-UV in HCT116 cells (Figure 2A). Additionally, closer inspection of SPUD splice junction coverage suggested the existence of several SPUD spliced variants (Figures 8B-D). Analysis of the number of reads split across each junction between exon 1 and SPUD-ALE (Figure 1C) revealed four predominant spliced isoforms (1–3, 1–4, 2–3, 2–4; Figure 8B), whereas only one junction between exons 2 and 3 was detected for *CDKN1A* full-length (Figure 8C). It is important to highlight that isoform 2-4 corresponded to the SPUD isoform characterized in the previous figures. Notably, the intronic sequence between exon 2 and 3 of *CDKN1A* full-length was almost undetectable in RNA-seq data (Figure 8C), whereas intron 1 appeared in poly(A)-selected RNA-seq (Figure 8B), suggesting it is more stable than CDKN1A downstream introns with slower splicing kinetics.

**Figure 8:**
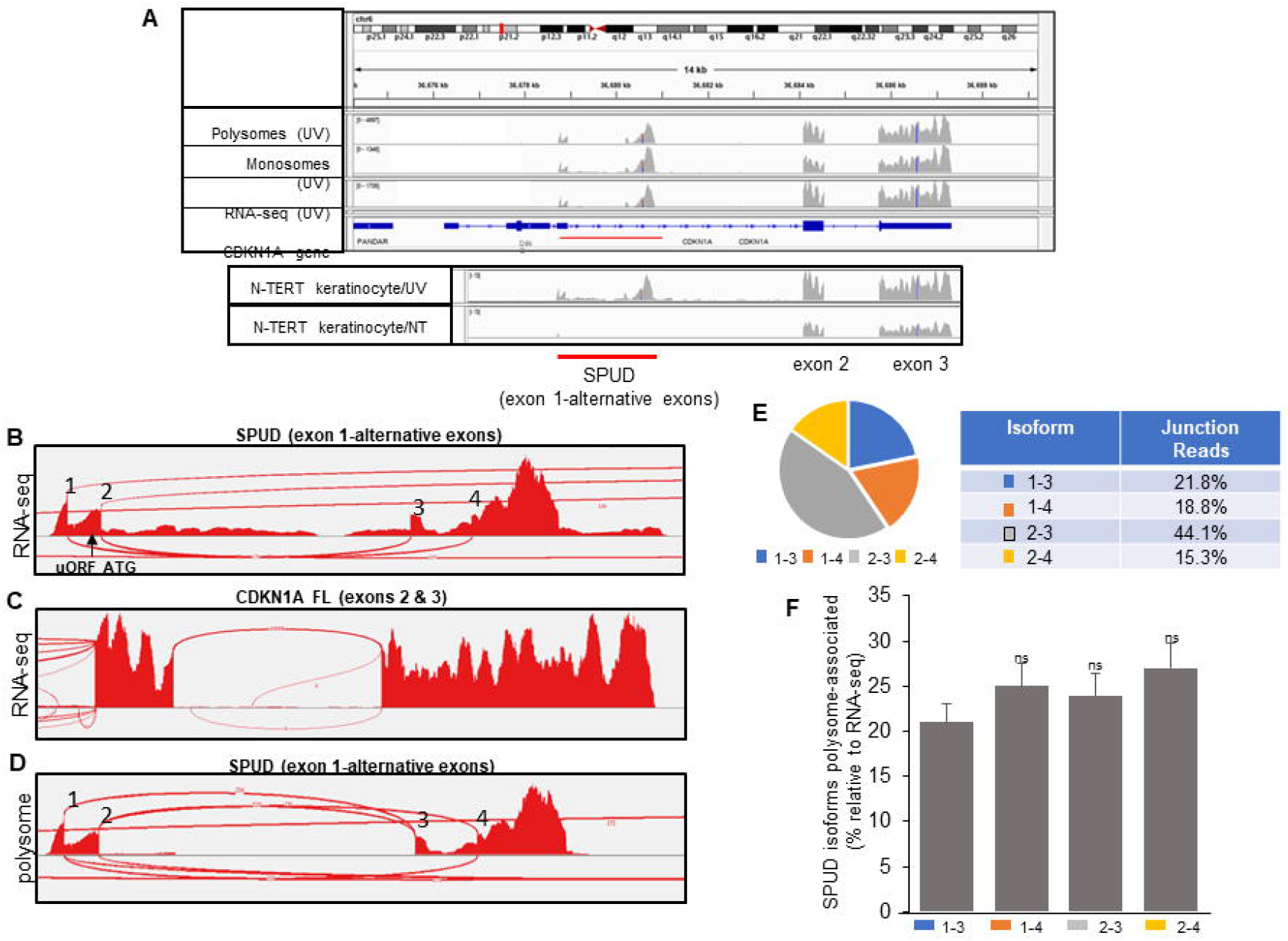
SPUD exists as multiple isoforms after UV treatment in keratinocytes. (**A**) High read coverage for SPUD-ALE RNA-seq analysis from total RNA, monosomes, and polysomes. Data obtained from N-TERT keratinocytes (GSE99745) before and after UV treatment were analyzed at CDKN1A gene using IGV viewer (hg38). NT, non-treated. The location of unspliced SPUD is indicated with a red line. (**B**) RNA-seq Sashimi plot for splice junction coverage of CDKN1A intron 1 upstream of APA event after UV treatment in N-TERT keratinocytes (GSE99745). The location of upstream ORF (uORF) ATG in SPUD mRNA is indicated. Each number corresponds to a utilized 5’ or 3’ splice site. (**C**) CDKN1A full-length mRNA utilizes only one set of splice sites. Sashimi plot showing one predominant splice junction for p21 mRNA coding exons. (**D**) All SPUD isoforms identified in (B) are detected associated with polysomes. Sashimi plot of RNA-seq from polysome fraction RNA showing the presence of all four isoforms. (**E**) Quantification of splice junction coverage for each isoform of SPUD using the annotations provided in (B). Percentage is the coverage as a proportion of the coverage for all four possible isoforms. Shown is the calculation from three biological replicates from N-TERT keratinocyte dataset (GSE99745). (**F**) All the SPUD isoforms associate equally with polysomes. Splice junction coverage percentage in the polysome fraction was divided by the average splice junction coverage percentage per isoform across three replicates of RNA-seq from N-TERT keratinocytes to provide a fraction of each isoforms’ polysome association relative to total RNA.

Investigation of polysome-associated RNAs (Figure 8D) showed the same four SPUD spliced isoforms detected by analysis of intronic reads present in spliced SPUD (Figure 8B); however, intronic reads present in unspliced SPUD were absent, consistent with the nuclear localization of this isoform (Figure 2E-F). Although the absolute proportion of each spliced isoform differed in different datasets (when comparing Figures 8E and 8F), the number of polysome-associated reads showed no enrichment of any isoform relative to RNA-seq reads (Figure 8F). Notably, only isoforms 2-3 and 2-4 contained the uORF ATG from CDKN1A mRNA (39,79). Neither isoform 1-3 or 1-4 contained ORFs >6aa, suggesting that SPUD may be ribosome-associated without participating in active translation for a specific ORF.

### RBPs CUGBP and CRT competitively bind both SPUD and CDKN1A full-length

Due to the apparent translational regulatory function of SPUD (Figures 6–7), as well as the finding that SPUD localizes to polysomes (Figure 8), we decided to investigate the potential functional overlapping of SPUD with other known translational regulators of p21. Most RBPs bind CDKN1A mRNA in the 3’UTR, with exception of CRT and CUGBP1 (80–81). CRT and CUGBP1 compete for binding exon 2 of CDKN1A mRNA and have opposing effects on p21 translation and on cell proliferation through p21 expression (80). While CUGBP1 activates p21 translation, CRT blocks translation by stabilizing of a stem–loop in CDKN1A full-length mRNA promoting a proliferative phenotype. Using *in vitro* biotinylated SPUD, RPD assays with HCT116 cell lysates showed that both CRT and CUGBP1 can bind SPUD (Figure 9A). Notably, an intramolecular stem-loop similar to the one described in exon 2 (80) can be predicted in SPUD-ALE and in exon 1 (Supplementary Figure S4A). The observation that p21 translational regulators can bind both CDKN1A full-length and SPUD transcripts further support the idea that the lncRNA SPUD might regulate p21 expression at the translational level. As with the CDKN1A full-length transcript (80), our RPD assays using limiting amounts of GST-CUGBP1 and increasing molar concentrations of GST-CRT showed that CRT and CUGBP1 competitively bind to SPUD (Figure 9B). Importantly, the binding of CRT and CUGBP1 to SPUD and CDKN1A changed during DDR progression (Figure 9C). RIP assays of samples from HCT116 cells exposed to UV treatment were IPed with antibodies against either CRT or CUGBP1. During early UV response (2 h), the p21 translational inhibitor CRT preferable bound SPUD over CDKN1A full-length and the p21 translational enhancer CUGBP1 preferable bound full-length over SPUD (Figure 9C). After 24 h, the binding of both proteins returned to the CDKN1A full-length/SPUD ratio observed in untreated cells. Importantly, the expression levels of both CRT and CUGBP1 did not change during the DDR progression (Supplementary Figure S4B), and the levels of CDKN1A isoforms returned to nontreated levels at 24 h after UV-treatment (Supplementary Figure S3A). Surprisingly, when the RIP assays were repeated after SPUD depletion, the binding of CRT to CDKN1A full-length increased both in stress and non-stress conditions (Figure 9D), suggestive of a competitive interaction of SPUD and full-length with CRT and providing a mechanism through which p21 translation is decreased.

**Figure 9:**
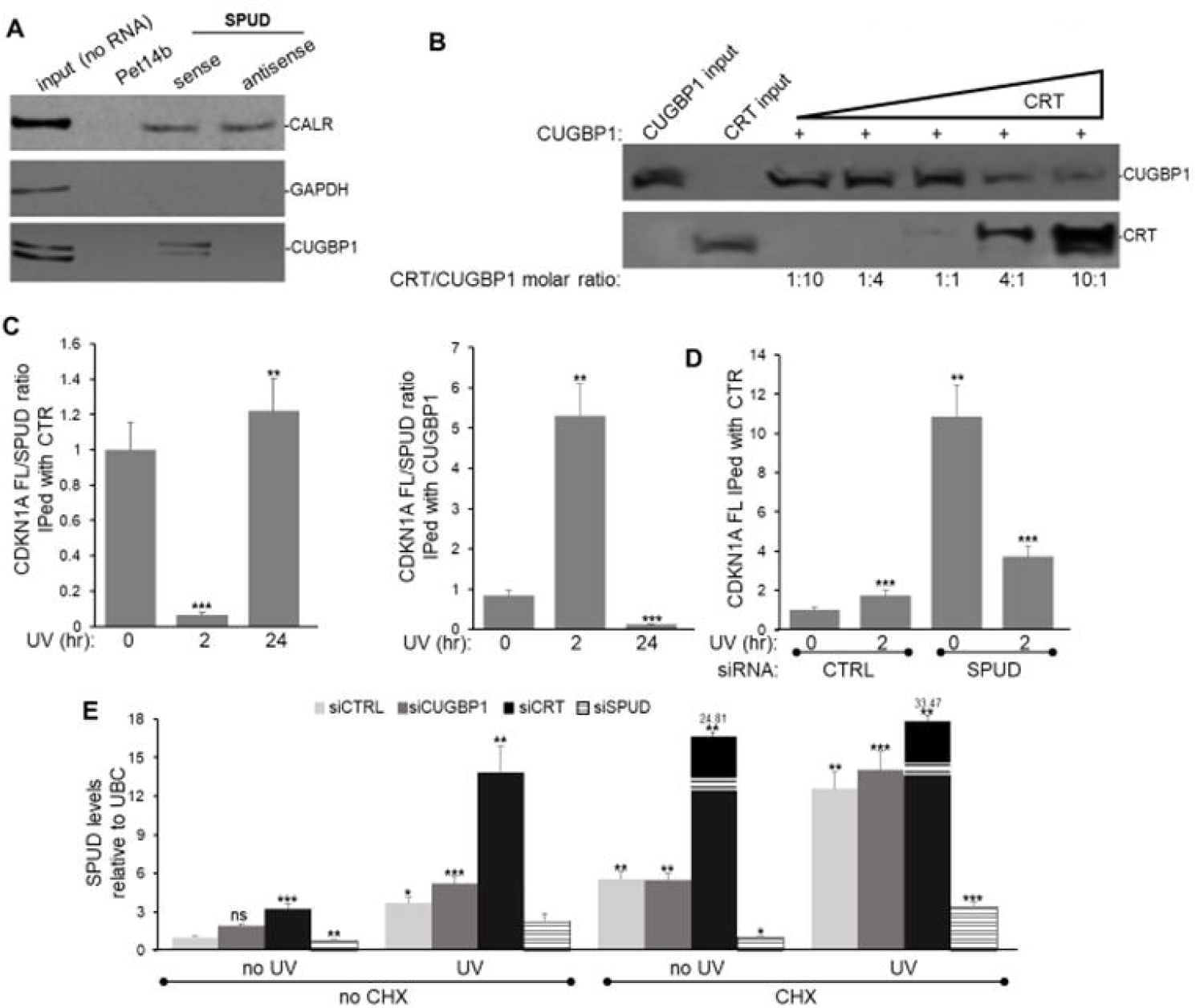
RBPs CUGBP and CRT bind directly to SPUD, and SPUD depletion results in an increase in CRT-CDKN1A full-length binding. (**A**) CUGBP and CRT bind directly to SPUD. Biotinylated sense and antisense SPUD were used in RPD assays with HCT116 cell lysate. Pet14b plasmid-generated RNA was used as a negative control. 5% input from HCT116 cell lysate is shown. As CRT binds a stem loop in exon 1, binding to sense and antisense is expected. CUGBP1 binds sequence specific, so only binding the sense is expected. GAPDH was used as loading control in Western blot analysis. A representative gel from 3 independent biological samples analyzed by triplicate is shown. (**B**) CUGBP and CRT compete for binding to *SPUD*. GST-CUGBP1 bound to biotinylated SPUD was mixed with increasing amounts of GST-CRT in RPD assays. Western blot analysis for the pulldown proteins is shown. A representative gel from 3 independent biological samples analyzed by triplicate is shown. (**C**) CRT binding to SPUD is favored over CDKN1A full-length binding at early times in the DDR, whereas CUGBP1 binding to CDKN1A full-length is favored over SPUD binding in non-treated cells and later in DDR. RNA-IP assay using either CRT, CUGBP1 or IgG antibody and samples from crosslinked HCT116 cells treated with UV irradiation (20 J/m^2^, recovery time indicated). Crosslinking was with 0.1% formaldehyde prior to sonication and immunoprecipitation. qRT-qPCR was performed on IP’ed transcripts. The CDKN1A/SPUD ratios observed in each condition were normalized to the values of non-treated samples. Data shown is from 3 independent biological samples analyzed by triplicate, SD (n=3).\*\**P*<0.001 and \*\*\**P*<0.0001. (**D**) RNA-IP performed as in (C) with samples of HCT116 cells depleted in SPUD by siSPUD treatment. Samples were analyzed as in (C). (**E**) UV and CHX treatment as well as CRT depletion increase *SPUD* transcript levels with minimal change on full-length CDKN1A levels. HCT116 cells were depleted in SPUD, CRT and CUGBP1 as in (D) and treated with either UV (20 J/m^2^), CHX (50 μg/ml) or CHX (2 μg/ml)/UV (20 J/m^2^) with 2 h recovery. qRT-qPCR was performed on RNA purified from cells treated in each condition. SPUD levels observed in each condition were normalized to the values of non-treated samples. Data shown is from 3 independent biological samples analyzed by triplicate, SD (n=3).\*\**P*<0.001 and \*\*\**P*<0.0001.

Consistent with SPUD’s role in controlling p21 translation, inhibition of translation by cycloheximide (CHX) treatment increased SPUD (~ 4-fold) to levels similar to those observed with UV treatment (2h, ~ 6-fold, Figure 9E). An additive effect was observed when cells were treated with both CHX and UV (~ 13-fold increase), which is abrogated by SPUD depletion. While CUGBP1 depletion did not affect SPUD levels, samples from CRT depleted cells showed a strong increase in SPUD levels in all four conditions (non-treated, CHX, UV and UV/CHX), indicative of potential feedback regulating full-length translation. These data suggest a potential translational mechanism to control p21 expression by which CTR, an inhibitor, binds SPUD in non-stress conditions keeping p21 levels low, whereas CUGBP1, an activator, binds the protein-coding mRNA and allows proper expression of p21 during DDR progression.

## Discussion

Cellular response to stress is achieved by the dynamic flux in gene expression. Post-transcriptional regulation of coding and non-coding RNA offers a fast method of adapting to a changing cellular environment. As intron-APA can alter the sequence composition of gene transcripts via intron retention and ALE inclusion (26), this alternative processing activation has the potential to diversify the functional output of genes or fine-tune the regulation of the canonical protein by the expression of additional proteins or non-coding RNAs (5). Therefore, understanding the complex interplay between APA events and genes involved in DDR/tumor suppression, such as *CDKN1A*/p21, can inform us about novel mechanisms to regulate cell-cycle checkpoints and the cell-fate decision between senescence and apoptosis during the progression of DDR. *CDKN1A* gene expression is highly regulated (81); early in DDR, despite the presence of p53 at the promoter, there is a delayed in p21 expression upon stresses like UVC or HU (17, 18), partially due to a block in transcription elongation somewhere in intron 1 (19,20,21,22). The fate of the truncated CDKN1A transcript generated in these conditions has not been investigated. In this study, we provided evidence that promoter-proximal intron-APA and alternative splicing are taking place in intron 1 of *CDKN1A*, generating an ALE in the SPUD transcript (Figures 1–2), under a variety of damaging conditions (Figures 5–6). *CDKN1A* intron 1 APA was first detected not only in our study of widespread APA occurrence in introns, biased towards the 5’ end of genes, after UV - treatment (32) but also in different dataset (Figure 1A). SPUD has low abundance compared to CDKN1A full-length isoform under basal conditions, is localized in the cytoplasm and highly stable, and induced in cancer and normal cells during DDR (Figure 2, 5). The RBP HuR binds to and promotes the stability of the unspliced precursor of SPUD, and the transcriptional repressor CTCF regulates SPUD levels (Figures 3–4). SPUD is a putative lncRNA that does not canonically encode for a protein and is regulated by p53, SPUD induction increases p21 but not CDKN1A full-length mRNA levels, affecting p21 functions in cell-cycle, CDK2 expression, and cell viability (Figures 6–7). Consistent with SPUD’s role on the induction of p21 expression at the translational level under stress conditions, SPUD is associated to polysomes (Figure 8). SPUD binding to the p21 translational activator CUGBP1 increases at early times after UV treatment and provides a mechanism for enhanced p21 translation following transient degradation of CDKN1A full-length transcript during S-phase and upon UV damage (82). Accordingly, SPUD depletion increases the level of CRT associated to CDKN1A full-length mRNA (Figure 9), probably resulting in the observed delay in p21 expression after UV treatment (17, 18). Taken together, our results indicate that intron-APA within *CDKN1A* produces a lncRNA that can fine tune the translation of p21 during DDR and cell-cycle.

Based on our results, we propose a model whereby intron-APA site activation and SPUD induction correlates with a decrease in U1 snRNA levels during DDR and is reversible by U1 snRNA overexpression (32). Importantly, studies have shown that inhibition of U1 function leads to activation of intron-APA events, resulting in shorter transcripts (30,83) and U1 snRNA overexpression mitigates UV-induced apoptosis. Consistent with this, UV-induced activation of PAS in intron 1 of CDKN1A occurs at times of UV-mediated depletion of U1 snRNA (~2-10 h after UV), and the suppression of APA and decrease of unspliced SPUD after UV occurs at times of recovery of U1 snRNA levels (>10 h after UV). The decrease in unspliced SPUD correlates with the increase of SPUD in progression of DDR, indicating that the precursor is spliced and exported to the cytoplasm, whereby the high levels of SPUD at later time points after UV treatment is reflective of its high stability. The delayed induction of CDKN1A full-length, but not of the lncRNA SPUD after UV treatment, might indicate that the proximal and canonical PAS can be recognized independently. However, we cannot discard the possibility that the proximal PAS can be activated sequentially following the canonical ones, suggesting that the same CDKN1A full-length transcript can undergo a multi-cleavage (84). Moreover, it has been shown that canonical PAS can enhance proximal-splice site usage (85). The delay in SPUD induction after UV compared with the unspliced SPUD induction indicates that APA occurs prior to 3’ splice site recognition. Interestingly, a measurable subset of SPUD precursor appears to remain unspliced and nuclear for an as-yet-unknown function.

Surprisingly, tandem PAS are in close proximity (~30 nt) within *CDKN1A* intron 1 (Figure 2A). Tandem PAS are usually located within the 3’UTR with a median distance between PAS of ~300 nt (86). Our 3’RACE results suggest that the first PAS encountered (ATTAAA) is preferentially used both before and after UV treatment (Figure 2D), whereas the downstream PAS (AATAAA) was used with less efficiency (and only after UV treatment). Although ATTAAA is the first encountered, it also has 5-6 times lower affinity for cleavage and polyadenylation specificity factor (CPSF) than AATAAA (87). However, it has also been shown that short (<10 bp) distances between PAS and GU-rich sequences might inhibit processing (88), explaining the lower usage of the second PAS. The region around the SPUD-ALE itself has been shown to be a central hub for binding activity of different factors; such as the transient binding of TGF-b-induced SMAD2/3 binding (89), MEF2C/D binding at SPUD-ALE (90), and p21-specific splicing regulator SKIP (64). The retention of multiple PAS and the functional association of different factors support the presence of these signals playing an important role.

The regulatory role of HuR on CDKN1A intron-APA and on the stability of the SPUD precursor is supported by previous studies that show the presence of HuR binding sites in the proximity of ALEs and APA signals by high-throughput and individual examples (91,92,93). The predicted HuR binding site in unspliced SPUD overlaps with a highly conserved region within CDKN1A intron 1; HuR binds specifically to the unspliced SPUD, but not the cytoplasmic transcript, both before and during the progression of UV-mediated DDR (Figure 4A-E). HuR’s regulatory role appears to be independent from the APA event itself, as HuR depletion under transcription inhibition conditions resulted in a decrease in the half-life of unspliced SPUD, but not SPUD, following UV induction (Figure 4F). SPUD levels did not change significantly by HuR depletion/actinomycin D treatment, which we attribute to the high stability of the transcript and suggesting that HuR does not play a role in splicing or SPUD stability.

PhyloCSF analysis of *CDKN1A* gene confirms the positive selection for CDKN1A full-length mRNA ORF but not for SPUD (Figure 6A), despite the conservation of a 3’ss, branchpoint and ATTAAA PAS in both rat and human. This supports the idea that SPUD is a lncRNA. This is further strengthened by the lack of *in vitro* translation of SPUD transcript or detection following expression of FLAG-tagged transcript beginning at exon 1 uORF ATG (Figure 6A-B). Furthermore, the multiple SPUD isoforms detected via existing RNA-seq possess similar polysome association, despite the presence of CDKN1A mRNA uORF in only a subset of isoforms (Figure 8F).

SPUD’s polysome association is consistent with the results showing that the lncRNA is a positive regulator of p21’s, but not p53’s, protein translation (Figure 6D). Polysome association of lncRNAs to regulate translation of other mRNAs has been observed previously, including through complementary base-pairing to short sequences (13,75,94). While several intergenic lncRNAs have been described in close proximity to the *CDKN1A* locus (15), including lincRNA-p21 that regulates CDKN1A *in cis* (12), SPUD is the first intragenic RNA *in trans* regulating p21 translation. We did not find any complementary sequences between SPUD and full-length transcript (13) between SPUD and CDKN1A full-length mRNA, rather we found that SPUD competes with fulllength mRNA for binding to two RBPs: the p21 translational repressor CRT and the p21 translational activator CUGBP1 (Figure 9A-B; 80). Loss of SPUD led to increased association of the full-length mRNA with CRT (Figure 9) and a reduction in p21 (Figure 7C) during DDR progression. Interestingly, CRT depletion increases the levels of SPUD (Figure 7D). While future experiments are necessary to test the details of this mechanism, it will be interesting to establish whether there are additional targets regulated at the translational by SPUD, particularly in a p21-null background, such in p21 -/- HCT116 cells, where SPUD expression is increased (Figure 5E) and a general dysregulation of protein expression and cellular functions have been described (67,68).

Despite the demonstration that only ~12% lncRNAs are expressed across all tissues (69), we have yet to observe a cell line that does not express SPUD (Figure 5A-C, Supplementary Figure S3B-C), most likely as it is transcribed intragenic to *CDKN1A*, which is ubiquitous. Rescuing transcription elongation of full-length also led to SPUD upregulation (Figure 6F). Consistent with this, both SPUD and CDKN1A full-length appear to utilize the same promoter in a p53-dependent manner (Figure 5C), supporting the idea that both SPUD and CDKN1A have the same transcription initiation.

Altogether, our data have provided evidence of a sense-strand intragenic lncRNA produced within CDKN1A that functions *in trans* to fine-tune p21 expression during DDR, reinforcing the functional interaction of lncRNAs and cell-cycle genes. Understanding the complex interplay between intron-APA events and canonical gene products involved in DDR/tumor suppression, such as SPUD/p21, might help us in identifying mechanisms that drive tumor progression, relapse, and treatment resistance.

## Data Availability

qRT-PCR was performed following MIQE Guidelines and details are included in Materials and Methods. siRNA off-target effects and overexpression data were included.

## Funding

This work was supported by National Cancer Institute, National Institutes of Health (NIH), R21 CA204610-01 and 1U54CA221704-01A (to FK).

## Acknowledgments

We thank Dr. Wilusz for CUGBP1-encoding plasmids, Dr. Michalak for CRT-encoding plasmids, Dr. B. Vogelstein for p21-null HCT116 cell line.

## Author contributions

Conceived and designed the experiments: MRM, AR, AD, and FEK. Performed the experiments: MRM, AR, AD, SV, DN, AY, SA, MM, MM, and GZ. Wrote and edited the paper: MRM, AR, AD, and FEK. Data Availability: The authors confirm that all data underlying the findings are fully available without restriction. All relevant data are within the paper.

## Conflict of interest

The authors declare no potential conflicts of interest.

## Supplemental Figure Legends

**Supplementary Figure 1:** Conserved elements within *CDKN1A* intron 1 corresponding to splicing and PAS. A) Complete RNA sequence for ENST00000462537 (SPUD). Highlighted in dark blue is the putative 3’ss. Highlighted in red are the tandem canonical PAS. B) Multi-vertebrate alignment in UCSC genome browser of region upstream of APA exon in CDKN1A intron 1. Light blue highlights are the putative conserved branchpoint(s) and 3’ splice site, respectively. Thin blue line is intronic region; thick blue line is exonic. C) Multi-vertebrate alignment in UCSC genome browser of the 3’ end of SPUD transcript in CDKN1A intron 1. Light blue highlights show the location and conservation of the ATTAAA and AATAAA polyadenylation signals. Exonic and intronic regions indicated as in (B).

**Supplementary Figure 2:** CTCF and HuR regulate CDKN1A isoforms and possess binding sites for both within CDKN1A intron 1. A) Sequence elements within SPUD highlighting HuR and CTCF binding sites. Shown is the entire unspliced SPUD transcript; yellow regions are exons, white regions are intronic. Sequence elements in green are putative HuR binding sites; pink are CTCF binding sites; teal are polyadenylation signals. B) Knockdown of HuR using siRNA. siRNA against HuR was transfected into HCT116 cells for 48 h followed by RNA extraction or 2 h UV treatment then extraction. Left, NEs were immunoblotted for HuR and LaminA as loading control. Right, quantification of immunoblot (n=3). C) siRNA against HuR leads to loss of CDKN1A isoforms. RNA extracted from HuR knockdown samples underwent semi-quantitative RT-PCR and agarose gel analysis for CDKN1A isoforms. UBC amplification was used as loading control; CDKN1A intron 2 was used as control for genomic DNA contamination. D) HuR overexpression does not impact CDKN1A isoforms. Mammalian expression vector containing HuR cDNA was transfected into HCT116 for 48 h followed by qRT-PCR analysis for indicated transcripts. Shown are the results from three independent assays. E) UCSC genome browser view of CDKN1A showing putative CTCF binding site from ENCODE ChIP-seq data in indicated cell lines. Peaks in red are UV-induced PASS reads. Leftmost red peak the position of SPUD APA event, Rightmost peak the full-length cleavage/polyadenylation site. F) HCT116 cells were transfected with siRNA against CTCF for 48 h followed by immunoblotting of NEs probing for CTCF and LaminA loading control. Shown is a representative gel from three replicates. Right, quantification of blot. ns, not significant; ** *P<0.001;* *** *P<0.0001*.

**Supplemental Figure 3.** A) SPUD overexpression leads to higher SPUD without affecting full-length mRNA levels. HCT116 cells were transfected with mammalian expression vector containing SPUD cDNA. After 48 h, cells were lysed and total RNA was extracted for analysis for CDKN1A isoforms. B) UV induces APA without inducing full-length in MCF7 breast cancer cells. Cultured cells were exposed to 20J/m^2^ UV and allowed to recover for 2 h. RNA was extracted and qRT-PCR analysis performed for the indicated isoforms (n=3). C) Overexpression of both unspliced SPUD and SPUD lead to upregulation of p21 protein levels. HCT116 cells were transfected with either empty FLAG-containing vector one of unspliced and spliced SPUD. Whole cell extracts were prepared and resolved by SDS-PAGE before probing for p21 or tubulin loading control (n=3). ns, not significant; ** *P<0.001*; **** *P<0.00001*.

**Supplemental Figure 4.** A) CUGBP and CRT levels are unchanged during UV-mediated DNA damage response. HCT116 cells were exposed to UV and allowed to recover for the described times prior to NE preparation. Samples were analyzed by Western blotting for the proteins indicated. B). Potential CUGBP-binding stem loops within SPUD. SPUD was analyzed using ViennaFold and stem-loops with the most consecutive base pairs were isolated. The location of either stem loop within SPUD is shown.

**Supplemental Figure 5.** Panel of oligonucleotides used in this study.

## Notes

### Competing Interest Statement

The authors have declared no competing interest.

