## Supplementary figures and images for "Long Non-Coding RNA Generated from *CDKN1A* Gene by Alternative Polyadenylation Regulates p21 Expression during DNA Damage Response"

### Figure 1 Supplementary

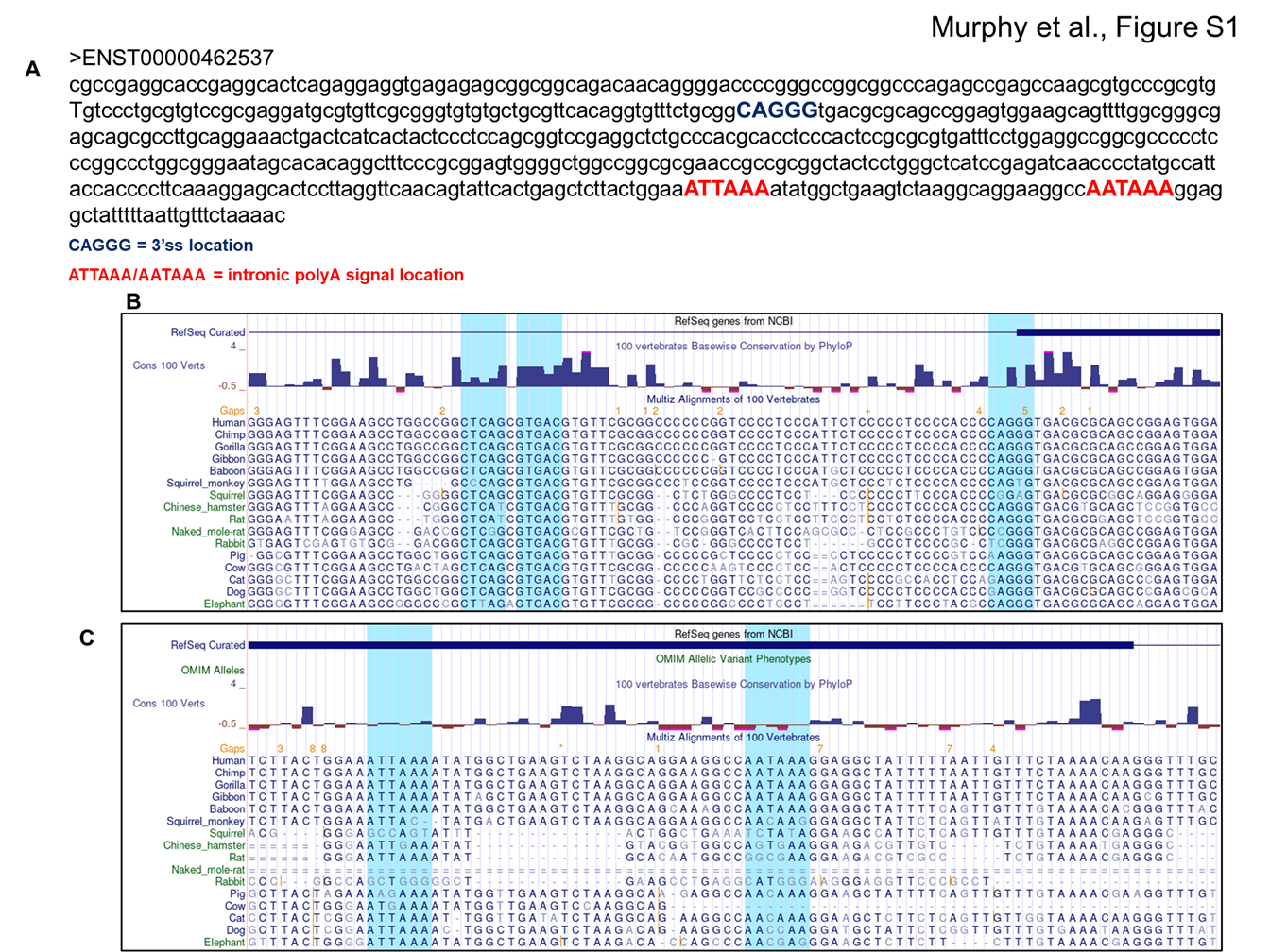

### Figure 2 Supplementary

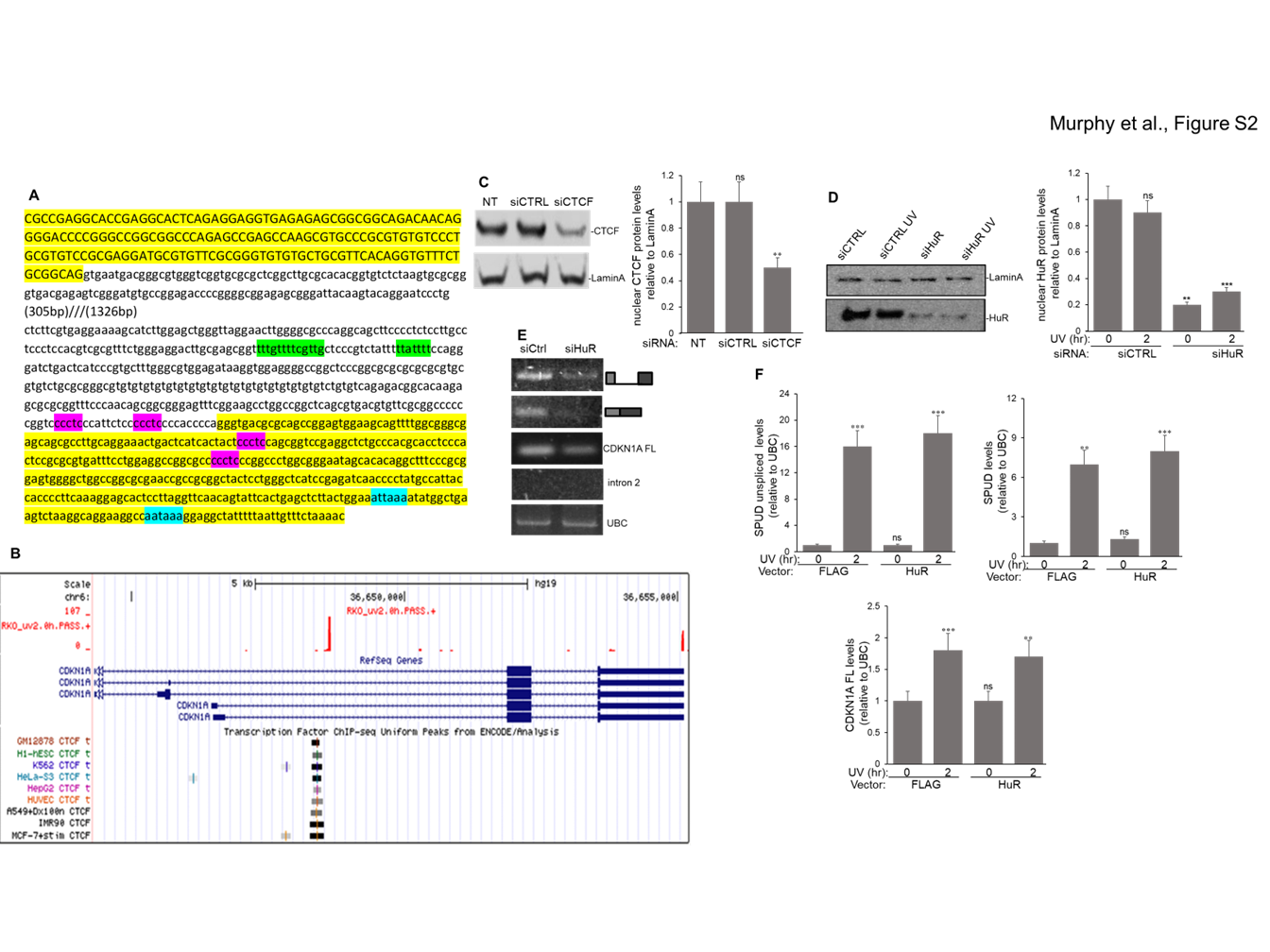

### Figure 3 Supplementary

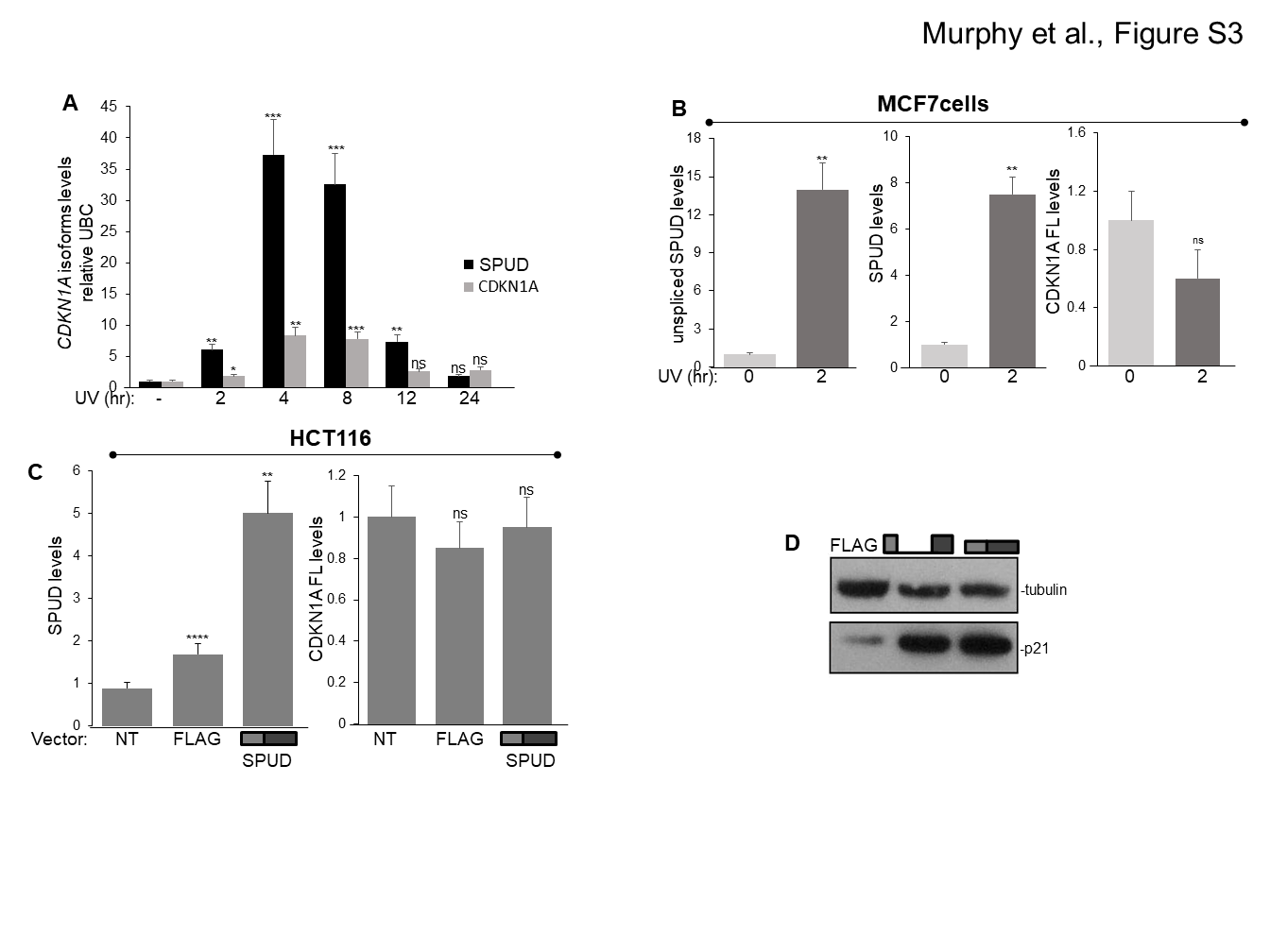

### Figure 4 Supplementary

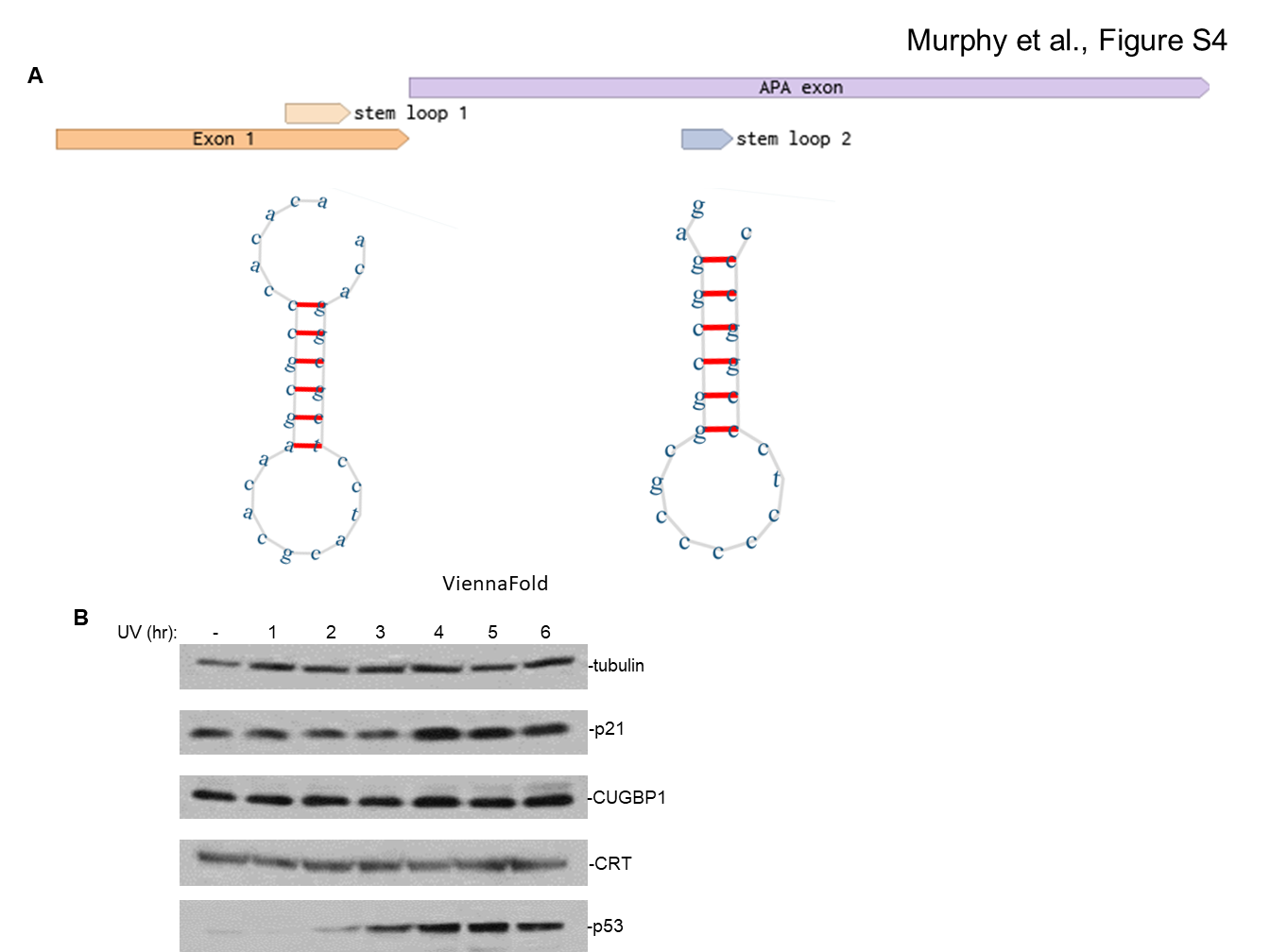

### Figure 5 Supplementary

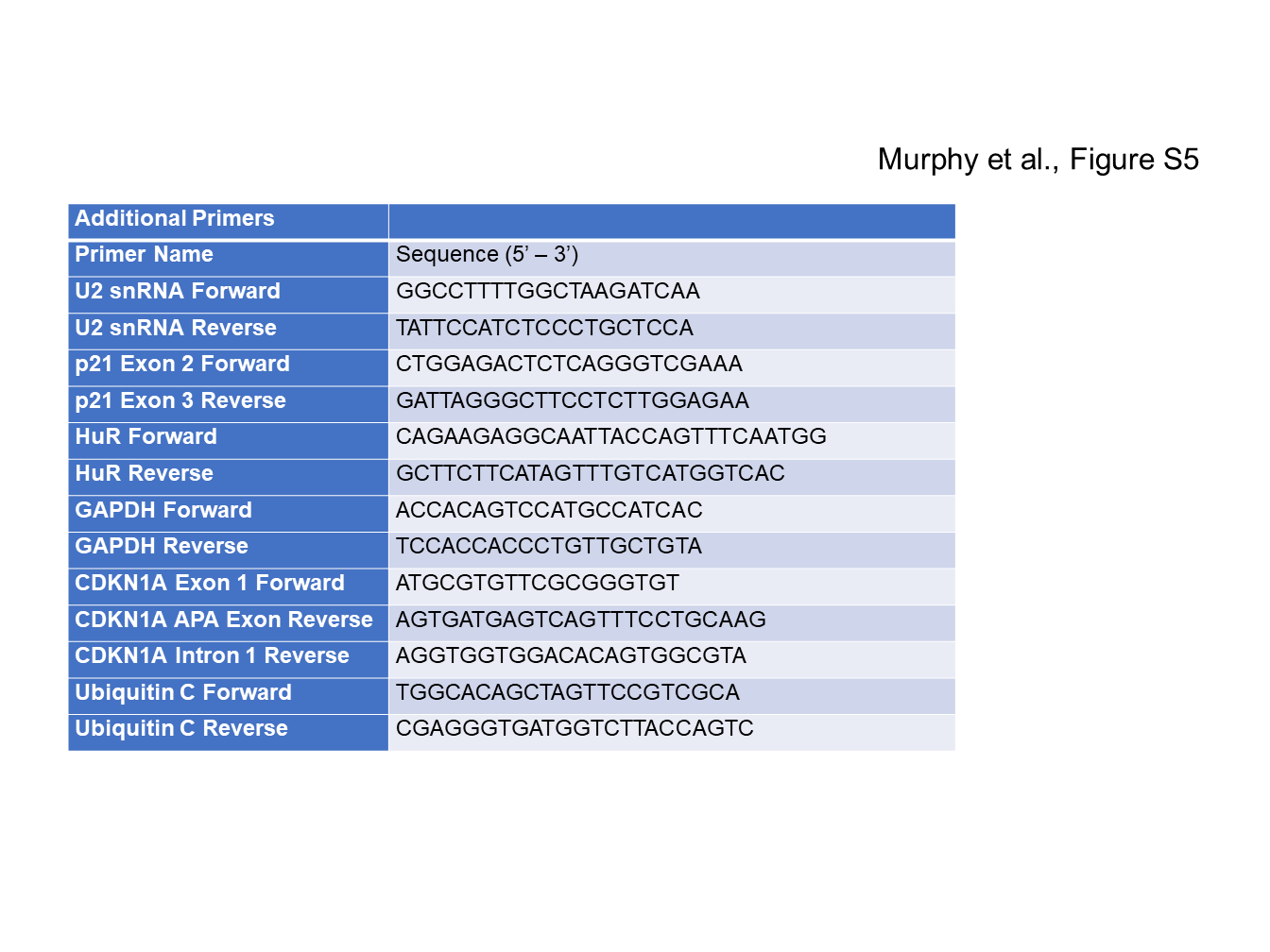
